# MetworkPy: A Python Package for Graph- and Information-theoretic Investigation of Metabolic Networks

**DOI:** 10.64898/2026.05.26.727944

**Authors:** Braden T. Griebel, Shuyi Ma

**Affiliations:** Center for Global Infectious Disease Research, Seattle Children’s Research Institute, Seattle, 98101, USA; Department of Chemical Engineering, University of Washington, Seattle, 98195, USA; Department of Pediatrics, University of Washington, Seattle, 98195, USA; Department of Global Health, University of Washington, Seattle, 98195, USA; Department of Microbiology, University of Washington, Seattle, 98195, USA; Viginia Merrill Bloedel Hearing Research Center, University of Washington, Seattle 98195, USA

**Keywords:** Genome-scale Metabolic Models, Graph Theory, Information Theory, Metabolic Modeling, Flux Balance Analysis, Flux Sampling

## Abstract

**Background:** Metabolism plays an important role in physiology and disease. *In silico* Genome-Scale Metabolic Models (GSMMs) are useful for informing analysis of metabolism and contextualizing high-throughput molecular profiling data. Flux balance analysis (FBA) based approaches can use GSMMs to predict metabolic reaction flux states and condition-specific alterations, but most FBA methods are limited by: 1) needing metabolic objective functions that are difficult to define experimentally, and 2) confining comparisons to extrema of reaction flux distributions.

**Results:** We introduce MetworkPy, a Python package that implements a flexible toolbox for objective-free graph- and information-theoretic analyses of GSMMs and flux distributions. MetworkPy includes methods to construct graph representations of GSMMs, to deconstruct GSMMs into topologically-aware gene sets that inform downstream bioinformatic analyses, and to assess shifts in flux distributions for groups of reactions. To demonstrate utility, we used MetworkPy to identify metabolic features perturbed by the transcription factor ArgR, a known regulator of arginine biosynthesis in *Mycobacterium tuberculosis*. MetworkPy successfully linked reaction flux shifts in ArgR’s transcriptome-constrained GSMM to arginine biosynthesis, which cannot be easily ascertained by FBA-based approaches.

**Conclusions:** MetworkPy is a customization-friendly toolkit for topological- and flux sampling based-analyses of GSMMs. MetworkPy’s tools provide multiple complementary metabolic contextualization approaches for microbial or eukaryotic genes of interest, and MetworkPy’s architecture facilitates user-designed workflows.

## BACKGROUND

Metabolism plays an important role in a broad spectrum of cellular responses, influencing disease susceptibility, immunity, and responsiveness to treatment [1–7]. Therefore, elucidating the metabolic context of cellular states can reveal mechanisms and inform intervention. A popular strategy to assess the metabolic context of cellular states is enrichment analysis, which compares a gene set of interest (e.g. derived from -omic profiling) against reference sets. These reference gene sets defined by gene ontology (GO) terms [8, 9] or canonical pathway databases (e.g. KEGG [10], Reactome [11, 12]) can aid metabolic contextualization and cross-species comparisons. However, they can lack condition-specific granularity, and they can be limited in their ability to assess enrichment of gene sets that cross canonically defined pathways. Genome-Scale Metabolic Models (GSMMs), which are *in-silico* descriptions of the metabolic reactions of an organism, have been used to contextualize -omic datasets and simulate metabolic states of cellular systems in condition-specific ways [13–17].

Flux balance analysis (FBA) and other constraint-based optimization approaches can use GSMMs to simulate metabolic states following chemical, genetic, regulatory, or environmental perturbations [18–22]. Such analyses have aided studies into metabolic regulation and metabolic responses during disease and therapy [14, 23–27]. However, most FBA-based methods are restricted by: 1) assumptions of optimality, which necessitate metabolic objective functions that are difficult to define experimentally, and 2) limitations in the number of possible reaction states evaluated during simulation to flux extrema [13, 17, 28].

To circumvent the need to define metabolic objective functions, approaches have been developed to investigate the full space of achievable reaction fluxes within a defined GSMM. The range of possible fluxes can be analyzed with flux variability analysis [29], which identifies the minimal and maximal fluxes for each reaction. However, the minimal and maximal fluxes do not provide information on which solutions within this range are feasible, nor which fluxes or combinations of fluxes are more likely [17]. In contrast, flux sampling provides a distribution of solutions throughout the feasible flux space [17, 30, 31]. Although sampling offers the promise of more comprehensive analysis, most methods for assessing the generated flux distribution analyze the marginal distribution of a single reaction at a time [17, 32–34], which loses some of the available higher dimensional information.

GSMMs can also offer valuable insights into metabolism through its representations of interactions between metabolic reactions, genes, and metabolites. Graph-theoretic approaches have been applied to systematically assess the topological properties of these interactions [35]. Graph-based approaches have aided prediction of gene essentiality [36–38] and informed studies linking metabolism with regulation [23] and disease comorbidity [39]. Network topology has also enabled analysis of metabolic modularity, highlighting the organization of the metabolic network into subunits and revealing how these subunits interact [40–42].

Despite the utility of analyzing metabolic topology, there is still no consensus on how to construct a graph from a GSMM [35]. Approaches include reaction and metabolite connectivity graphs, as well as bipartite graphs with nodes representing both reactions and metabolites [23, 35, 43–45]. An additional challenge lies in dealing with currency or pool metabolites, such as adenosine triphosphate (ATP) or water (H_2_O), whose high connectivity may distort the topological properties of the graphs [35]. Approaches to mitigate these issues include removing highly connected metabolites [23, 46, 47], manual removal of metabolites from reactions where they act as carriers [42], automatic removal of currency metabolite pairs [48], and using flux weighting to mitigate the influence of these metabolites [45].

To empower topological and flux sampling-based analyses of GSMMs, we have developed the Python package MetworkPy. MetworkPy combines customizable methods for creating and analyzing network and subnetwork representations of GSMMs (**Figure 1**) with methods for generating and analyzing flux sampling distributions of reactions and pathways (**Figure 2**). MetworkPy further builds on the foundations of this toolkit with several analysis methods designed to contextualize gene sets within GSMMs; these analyses complement commonly used enrichment approaches. MetworkPy’s ability to create and evaluate a range of different graph and flux sampling representations of GSMMs not only fosters user-driven tailoring of analyses to suit custom needs, but also enhances robustness by enabling multiple alternative analysis representations to augment each other.

**Figure 1.**
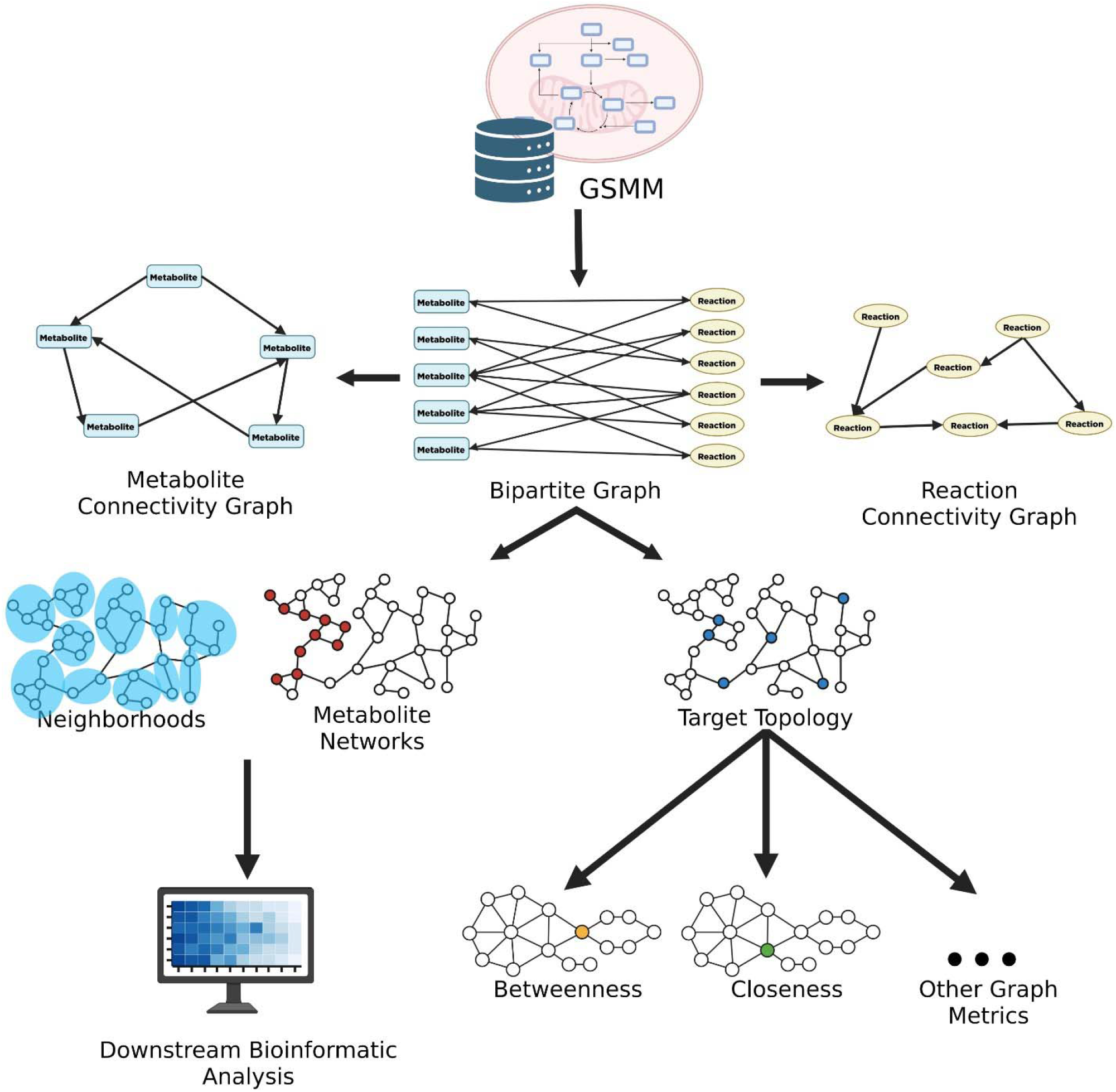
Overview of MetworkPy’s topology analysis pipeline. *MetworkPy converts an input GSMM into a bipartite graph representation* of stoichiometric connectivity featuring both reactions and metabolites *(with optional weights). This bipartite network can be used to analyze metabolic topology or to identify metabolic subsets for downstream analysis*. The bipartite network can *be optionally projected onto* only *reactions or metabolites* before executing downstream analysis. *Created in BioRender. Griebel, B. (2027)* https://BioRender.com/5sgzvxr.

**Figure 2.**
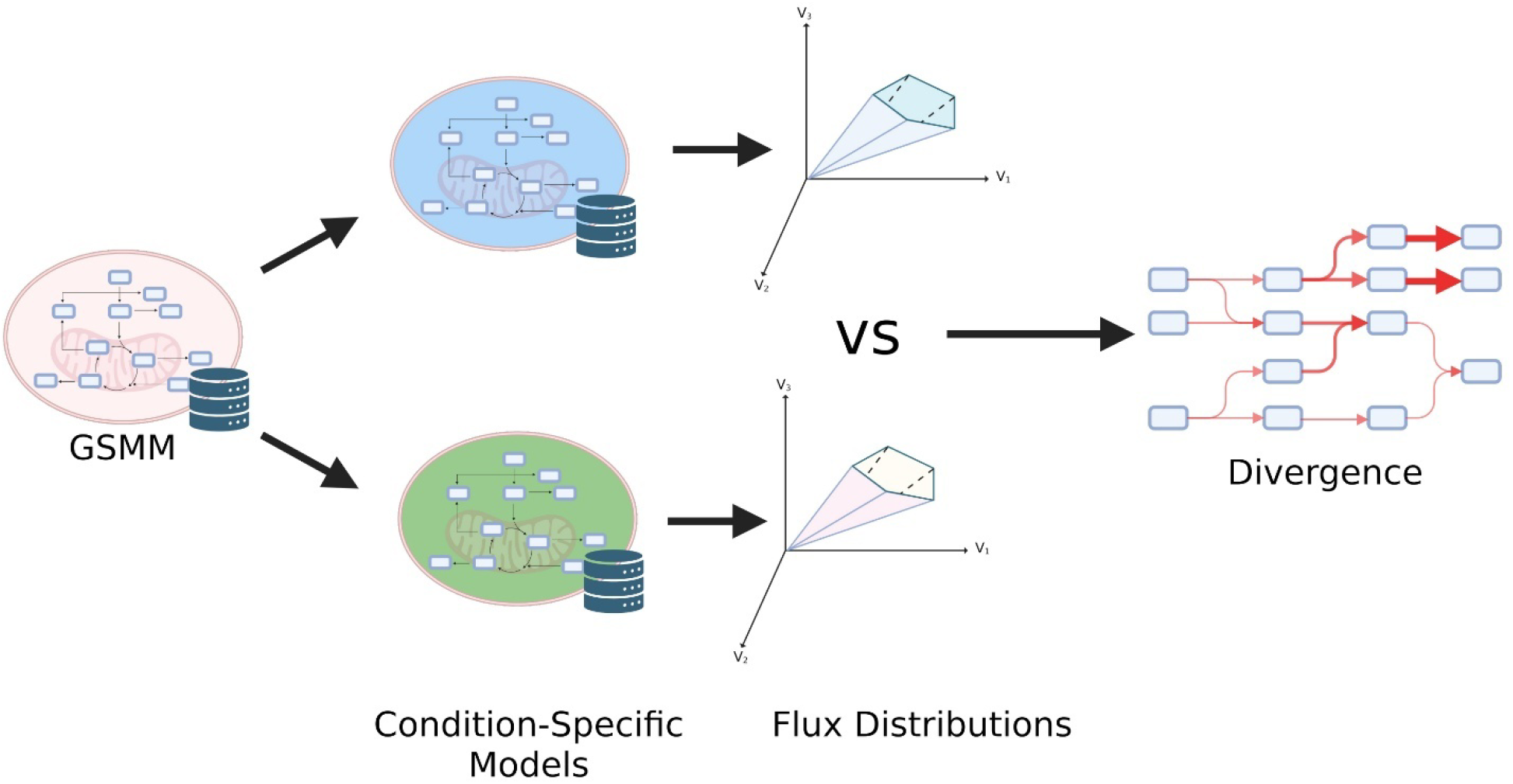
Overview of Flux Sampling Divergence Analysis (FSDA). Two condition-specific metabolic models (generated by constraining a base GSMM with gene expression or other - omic data, or by imposing another model manipulation like a simulated knockout) are flux sampled. The resulting distributions of fluxes for reactions or groups of reactions are then compared using Kullback-Leibler or Jensen-Shannon divergence metrics. Created in BioRender. Griebel, B. (2027) https://BioRender.com/gcewf00.

## IMPLEMENTATION

### Package Overview

MetworkPy offers a flexible toolkit for creating and analyzing graph and subgraph representations of an input GSMM (**Figure 1**). These network representations are in the form of NetworkX Graphs (or DiGraphs), or SciPy sparse matrices (currently only the bipartite network). These networks can be deconstructed using MetworkPy into neighborhoods and metabolite subnetworks, and the genes, reactions, or metabolites represented in these subnetworks can be used as metabolism-informed reference sets for enrichment analysis or other downstream bioinformatic pipelines. MetworkPy uses these network deconstruction methods, along with divergence-based analysis of flux sampling distributions (**Figure 2**), to contextualize gene sets of interest within the GSMM.

MetworkPy accepts as input: one or more “target” gene sets of interest and a GSMM. The input target gene sets can be derived from a broad range of bioinformatic workflows, such as differential gene or protein expression analysis, post-translational modification identification, or mutation analysis. To enable condition-specific granularity of analysis outputs, MetworkPy includes an implementation of the iMAT method [18, 19] to constrain input GSMMs with transcriptomic data.

MetworkPy can be used as a library for software developed in Python, as part of gene set-oriented bioinformatic pipelines, or in an interactive Python environment (i.e. Jupyter or Marimo). Written in pure Python, MetworkPy can also be easily modified. A summary of methods and functions featured in MetworkPy is described in **Tables S1 and S2** in **Additional File 1**. To illustrate how MetworkPy’s analyses can assess various metabolic features, we have applied MetworkPy’s main analysis workflows to a small simulation metabolic model (see **Additional File 1 S2.1** for description of the model and **Additional File 2** for analysis results).

### Metabolic Connectivity Network Generation

To facilitate analyses that leverage metabolic topology, MetworkPy offers flexible methods for converting a GSMM into a graph representation (**Figure 1**). A GSMM is first converted into a ***stoichiometric connectivity graph***, which we represent as a bipartite network with nodes representing both reactions and metabolites. Edges connect reactions to metabolites (and metabolites to reactions) if a metabolite participates in a reaction (See **S2.2.1** and **Figure S2 in Additional File 1**, and **Additional File 2**). Representing the GSMM as a bipartite graph allows for independent weighting and directionality of edges between a reaction and its metabolites. This weighting is flexible, allowing for user-defined forward and reverse weights for each reaction. Weights can be assigned based on stoichiometric coefficients, input -omics measurements, fluxes simulated via parsimonious flux balance analysis (pFBA) [49], geometric flux balance analysis (gFBA) [50], or flux variability analysis (FVA) [29]. Reactions weights can optionally be multiplied by the metabolite stoichiometric coefficients, and the production and reaction weights can be reweighted by a user-designated function if desired. The graph creation methods of MetworkPy also include several strategies for handling currency and/or carrier metabolites to avoid distortion of the topological properties of the graph (See **Additional File 1 S2.2.1)**.

Sometimes, it is desirable to work with a graph that includes only reaction or only metabolite nodes. MetworkPy includes methods for projecting the bipartite metabolic network onto reaction or metabolite nodes. These methods allow for a variety of projection algorithms that enable flexible handling of edge weights (see **Additional File 1 S2.2.1**). MetworkPy also includes helper functions to create these networks directly from the GSMM. The reaction connectivity network can be further converted into a gene connectivity network (where genes are connected if they are associated with the same, or neighboring, reactions) to enable topology-aware analyses directly centering metabolic genes.

If there are multiple groups of nodes of interest in one of these graph representations, MetworkPy can also facilitate the creation of a target set connectivity graph (inspired by [39]), where nodes represent target sets of nodes from the original graph and are connected if the node target sets neighbor each other in the original graph (see **Additional File 1 S2.2.1**).

All metabolic topology graphs are represented as NetworkX Graphs [51]. This enables calculation of established graph-theoretic properties with pre-existing tools. Key graph properties assessed include centrality and community detection [51, 52] (see **Additional File 1 S2.3.1** and **S3.3**).

### Network-centric Gene Enrichment Analysis

Examining activity of metabolic pathways or subnetworks increases analysis robustness, since consistent activity over a set of related reactions is less likely to arise by chance. The highly interconnected nature of metabolism means that analyzing canonically defined pathways as separate entities, as done in conventional pathway enrichment analyses, can obscure the relationships between them. To address this challenge, MetworkPy defines graph-theoretic subnetworks from stoichiometric connectivity graphs. MetworkPy’s topology-driven subnetworks can highlight changes in reaction sets that are proximally connected in the metabolic network at the genome scale, even if they are not conventionally represented as linked in canonically described pathways. MetworkPy’s graph-theoretic strategy therefore has the advantage of identifying emergent metabolic associations that might arise in non-model organisms or systems that might be missed with conventional pathway definitions.

MetworkPy implements two types of subnetworks: ***reaction neighborhoods*** (described in **Additional File 1 S2.2.2**) and ***metabolite subnetworks*** (similar to those used in [24], described in **Additional File 1 S2.2.3**). These subnetworks can be used to inform network overlap analyses described further below or exported as gene set inputs to other bioinformatic pipelines that accept a customized gene set input (e.g., GSEA [53], GSVA [54], etc.).

Reaction neighborhoods are sets of reactions (or associated genes) that are proximally connected in the stoichiometric connectivity graph. Previous work has shown that neighboring reactions or metabolites within the connectivity network are informative about metabolic alterations [43] and their consequences on disease physiology [39]. MetworkPy generalizes neighbors in the connectivity network to include reactions within a user-configurable distance. MetworkPy offers several methods of working with these neighborhoods, including iterators over node or gene neighborhoods in a graph, and higher-order functions for mapping a function over these neighborhoods (in parallel). Examples of the utility of these neighborhoods are analysis using the network density metric and fuzzy intersection methods, described below.

Metabolite subnetworks identify the reactions and associated genes in the GSMM that are required to synthesize, or which consume each metabolite. Metabolite subnetwork definitions facilitate direct interrogation of how perturbations throughout a GSMM might influence metabolite levels. Differential Producibility Analysis (DPA), developed by [24], uses metabolite synthesis networks to identify metabolic alterations in the bacterial pathogen *Mycobacterium tuberculosis* (Mtb) [24, 25]. MetworkPy identifies the reactions in metabolite synthesis networks in a way similar to DPA [24], and extends the functionality by also defining subnetworks of reactions and associated genes that consume each metabolite.

Assessing the overlap of multiple sets of target genes can highlight neighborhoods or subnetworks that are concomitantly impacted by these different targets. Though enrichment metrics can be used for overlap analysis, they underreport instances of overlap when there are many genes in the target sets outside the overlapping region. Overlaps can also be obscured if there are no exact matches between the gene sets of interest. To overcome these challenges, MetworkPy establishes a novel ***density*** metric for overlap analysis. Density represents the proportion of genes or reactions in a reaction neighborhood or metabolite subnetwork that are included in a target set of interest. Given that density is assessed in a particular subnetwork, it is not impacted by genes in other subnetworks; thus, overlap evaluation is not diluted by out-of-scope genes. Furthermore, density converts the gene set into a fuzzy [55] reaction set (MetworkPy also includes alternate approaches for converting a gene set into a fuzzy reaction set, see **Additional File 1 S2.3.2**). When combined with fuzzy intersection analysis methods (for which MetworkPy includes several options, including one based on robust rank aggregation [56]), density can identify instances of fuzzy overlap amongst multiple gene sets (see **Additional File 1 S2.3.2**). Although MetworkPy’s density metric is geared toward analyzing GSMMs, these MetworkPy methods can also be applied to other biological networks.

In addition to density and enrichment analysis, MetworkPy also offers methods for clustering target sets of nodes within the stoichiometric connectivity graphs. The distance between target node sets can be returned as a matrix to facilitate clustering with methods that accept precomputed distances (e.g. from scikit-learn [57]). MetworkPy includes methods for agglomerative hierarchical clustering of node groups in the metabolic network, which can optionally return a linkage matrix that can be used with SciPy [58] to create a dendrogram (see **Additional File 1 S2.3.4**).

### Information-theoretic Reaction Flux Divergence Analysis

Comparative FBA methods that investigate metabolic perturbations in GSMMs often focus on a single or a small number of feasible solutions (as in FVA, pFBA, etc.). This can miss a large portion of the flux space. Metabolic reaction flux sampling can assess distributions of feasible reaction flux solutions more comprehensively. MetworkPy includes an implementation of the corner sampling approach [59] and can also leverage flux distributions calculated from other sampling approaches (e.g., artificial centering hit-and-run (ACHR) [60] and optGp [30]) to assess the entire feasible flux space.

Results of flux sampling simulations have typically been investigated using statistical tests comparing the marginal distributions of individual reactions [17, 33, 34]. While this approach can identify metabolic alterations, it is limited in two key respects: 1) this approach is designed to compare a single reaction’s marginal flux distribution at a time, so changes in groups of reactions cannot be evaluated; and 2) this approach is designed to assess changes in a specific summary statistic. An alternative strategy to address these limitations is to use a statistical distance measure, such as the Kullback-Leibler divergence [59].

MetworkPy expands the use of divergence for investigating the feasible flux space [59, 61, 62] by including implementations of Kullback-Leibler [63] and Jensen-Shannon [64] divergence measures, based on the approaches of [65, 66], which can handle higher-dimensional distributions (**Figure 2**, see **Additional File 1 S2.4** for details). MetworkPy’s Flux Sampling Divergence Analysis (FSDA) offers two key advantages: 1) divergence can evaluate changes for sets of reactions, making it compatible with assessing changes in pathways, metabolic subnetworks, and/or user-configurable sets of reactions; and 2) divergence is sensitive to a wide range of changes to the distribution, including shifts in mean or variance.

## RESULTS AND DISCUSSION

### ArgR Target Topology Analysis

To demonstrate the utility of MetworkPy’s topology-aware metabolic contextualization of target gene sets we applied these analyses to assess the metabolic regulation of the *Mycobacterium tuberculosis* (Mtb) transcription factor ArgR (Rv1657), a known regulator of arginine biosynthesis in Mtb [67–69].

We started by using MetworkPy to identify metabolite subnetworks and reaction neighborhoods in the Mtb GSMM iEK1011_v2 [16] (**Additional File 1 S3.2** for details). Then, we evaluated the enrichment of genes targeted by ArgR (defined here as genes differentially expressed in an Mtb strain that recombinantly overexpressed *argR* [69]) in these subnetworks. As expected, we found increased density and significant enrichment for the regulatory target genes of ArgR in the arginine metabolite subnetwork (Benjamini-Hochberg false discovery rate (FDR) adjusted [70] enrichment p = 1.86*10^-11^, ranked as most significant) and the reaction neighborhoods around reactions in arginine biosynthesis: acetylglutamate kinase (ACGK, FDR adjusted enrichment p = 9.1*10^-4^, ranked as most significant) and ornithine carbamyltransferase (OCBT, FDR adjusted enrichment p = 1.8*10^-2^, ranked as second most significant) (see **Figure 3**, **Additional File 1 S3.2**, and **Additional File 3**).

**Figure 3.**
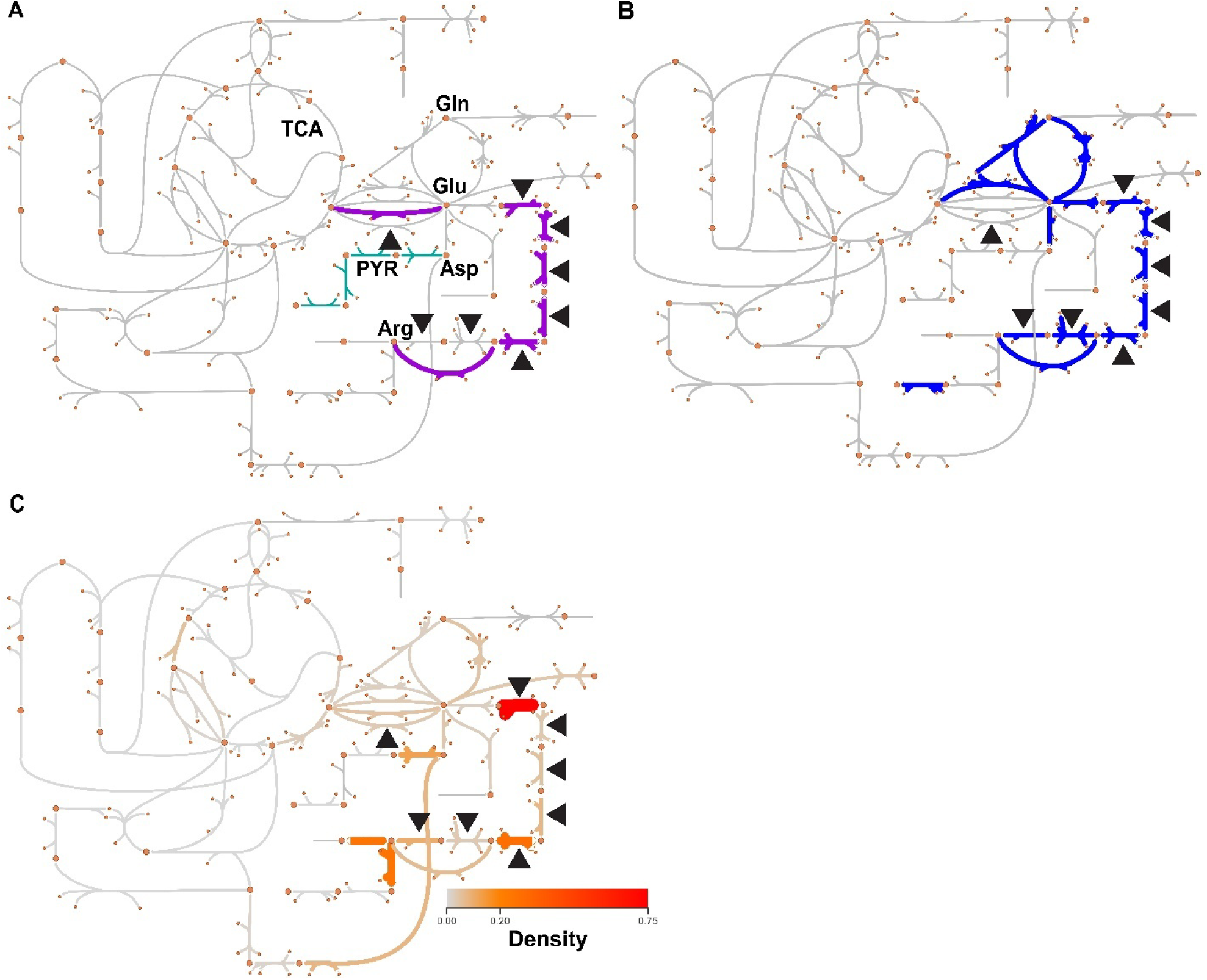
ArgR target density and arginine biosynthesis pathways. Each panel shows an Escher Map [72] of nitrogen and central carbon metabolism, marked with information on arginine biosynthesis and ArgR-associated reactions. Both large and small circles indicate metabolites, and the connecting curved or straight lines indicate reactions (see **Additional File 1 Figure S11** for a version of the Escher Map that includes reaction and metabolite labels). The black triangles indicate the reactions associated with genes targeted by ArgR based on data from [69]. **(A)** Shows reactions (highlighted in purple) that are annotated in the iEK1011_v2 metabolic model as being part of the ‘Arginine and Proline Metabolism’ subsystem [16]. Reactions in the ‘Purine and Pyrimidine Biosynthesis’ subsystem are colored teal. Specific metabolites and pathways of interest are labeled: PYR – Purine and Pyrimidine Biosynthesis, TCA - Tricarboxylic acid cycle, Arg - Arginine, Asp - Aspartate, Glu - Glutamate, Gln - Glutamine. **(B)** KEGG’s ‘Arginine biosynthesis’ pathway [10] is highlighted in dark blue. **(C)** ArgR target density values calculated with MetworkPy (based on the aforementioned targets [69] defined by the black triangles) are mapped onto the Escher Map; the color and thickness of the reactions are both proportional to the density level.

Second, by applying MetworkPy’s density metric, we detected higher density levels for reaction neighborhoods proximate to arginine metabolism, including reactions associated with purine and pyrimidine metabolism (**Figure 3**). Notably, associations with purine and pyrimidine metabolism were not detectable using GSEA with KEGG pathways or reaction subsystems defined by iEK1011_v2 [16] (see **Additional File 1 S3.2** and **Additional File 3**).

Third, we analyzed the topology of ArgR targets using MetworkPy’s subset centrality methods. These methods similarly indicated the relationship between ArgR and arginine biosynthesis-associated metabolites and reactions (see **Additional File 1 S3.3** and **Additional File 3**).

As a demonstration of integrating the gene sets identified by MetworkPy with other bioinformatic analyses, we combined the identified metabolite synthesis subnetworks with Gene Set Variability Analysis (GSVA) [54] as implemented in the decoupler package [71]. Using the metabolite subnetworks as gene set inputs to GSVA, we analyzed the transcriptomes elicited in the aforementioned Mtb strain that recombinantly overexpressed *argR* [69]. This approach identified arginine and closely related metabolites (e.g., ornithine) as the most altered upon ArgR overexpression (see **Additional File 1 S3.2** and **Additional File 3**).

### Gene Knock-out Divergence

A powerful application of MetworkPy’s FSDA is its ability to quantify network-level metabolic flux impacts of disrupting either individual or multiple metabolic genes (defined by user input or molecular profiling datasets). To demonstrate this, we examined the flux divergence in metabolite synthesis networks for all single gene knockouts in the iEK1011_v2 model of Mtb. Knockout simulations of essential genes (profiled by [73]; data retrieved from [74]) elicited significant divergence in the biomass reaction (Mann-Whitney U-test p = 4.5*10^-66^). Further, the divergence of biomass reaction is also significantly correlated with the Vulnerability Index of [74] (Pearson R = -0.61; p = 1.69*10^-103^). Using the divergence of the biomass, gene essentiality could be predicted with an AUC-ROC = 0.84, virtually identical to the AUC-ROC achieved by the traditional FBA optimal growth analysis (AUC-ROC = 0.84).

MetworkPy’s FSDA approach can yield information on more than just the biomass reaction. FSDA can also be used to evaluate impacts on both individual reactions and pathways, reaction neighborhoods, and/or metabolite production/consumption outcomes. For example, MetworkPy showed that genes targeted by ArgR caused significantly higher divergence values in arginine biosynthesis following knockout than those not targeted by ArgR (FDR adjusted Mann-Whitney U-test p = 2.23*10^-6^; see **Additional File 1 S3.1** and **Additional File 3**).

### Condition-specific GSMM Divergence

When paired with condition-specific GSMMs (such as models defined by gene expression patterns profiled under different experimental conditions), MetworkPy’s FSDA highlights and ranks reactions, pathways, reaction neighborhoods, or metabolite subnetworks with significantly altered flux distributions between the conditional models. To demonstrate utility, we compared the metabolic consequences of overexpressing ArgR relative to overexpressing each of the other Mtb TFs. We based this analysis on gene expression data from [69], generated for a library of Mtb strains recombinantly engineered to overexpress TFs (here referred to as TFOE strains). We used MetworkPy’s implementation of iMAT [18, 19] (see **Additional File 1 S2.5**) to generate condition-specific GSMMs for each TFOE strain, and to generate a reference GSMM using the median expression across all measured TFOE strains. We evaluated the divergence between each TFOE-specific GSMM and the median model for all individual reactions, metabolite synthesis subnetworks, model subsystems, and KEGG pathways. The divergence values were converted into z-scores for each network by normalizing across the TFs. For ArgR, the most perturbed metabolite synthesis network is ornithine, a metabolic precursor of arginine. Additionally, of the top 10 most perturbed reactions, 7 are in the “Arginine and Proline Metabolism” subsystem of the iEK1011_v2 model, with the other 3 being in the closely related “Alanine, Aspartate, and Glutamate Metabolism” subsystem.

We also combined the unnormalized divergence values with the neighborhood enrichment measures using Robust Rank Aggregation (RRA)[56]. We found that of the top 10 ranked reactions, 6 were in “Arginine and Proline metabolism”, and another 3 were in the closely related “Alanine, Aspartate, and Glutamate Metabolism” (see **Additional File 1 S3.5** for more details).

In contrast, in comparing the pFBA-simulated fluxes generated from the ArgR iMAT model relative to the pFBA fluxes [49] from the median iMAT model, the top perturbed reactions are associated with the “Transport”, “Oxidative Phosphorylation”, “Redox Metabolism”, “Extracellular Exchange”, and “Purine and Pyrimidine Biosynthesis” subsystems (see **Additional File 1 S3.4** and **Additional File 3**). We further compared alternative flux sampling-based approaches, evaluating differences in flux samples taken from the median iMAT model versus those taken from the ArgR iMAT model. We compared the marginal distributions for the reactions using several statistical tests (Student’s t-test, Mann–Whitney U test, Kolmogorov–Smirnov test, Kruskal–Wallis test). Though these tests were able to identify perturbations in reaction in the arginine biosynthesis pathway, none were able to differentiate these arginine reactions from reactions in the rest of metabolism, ranging from ‘Glycolysis/Gluconeogenesis’ to ‘Nucleotide Salvage Pathway’ (see **Additional File 1 S3.4**, and **Additional File 3**).

### Software Comparison

There are a variety of other applications and libraries that overlap in functionality with subsets of MetworkPy’s utilities. In our literature review, we have found three classes of analyses that have overlap: 1) metabolic topology analysis, 2) flux sampling analysis, and 3) metabolite subnetwork analysis. We will describe comparisons against each class in the following sections. Overall, we find that MetworkPy brings together and generalizes diverse analysis approaches, providing a conveniently integrated library.

#### Metabolic topology analysis

The topological properties of metabolic networks have been analyzed in a variety of other studies (e.g., [36–38, 40, 42–47, 75–80]). However, in many cases the code for generating and analyzing the metabolic graph from the GSMM is not always available [36–38, 40, 43, 47, 75–77], or is deprecated [46, 78], or the method of constructing the metabolic network is specific to the analysis pipeline being developed [42, 44, 45, 79, 80].

Fluxer [81] and MetExplore [82, 83] are web applications that can be used to visualize and explore metabolic networks. Fluxer and MetExplore are graphical user interface (GUI) tools. GUIs simplify usage; however, GUIs can restrict flexibility and make automation and integration as part of a bioinformatic pipeline more challenging. MetworkPy focuses on usage as a software library, allowing for more flexibility in constructing graph representations from GSMMs, and simpler integration with pre-existing Python graph analysis tools (via NetworkX and SciPy). MetworkPy also offers additional tools for analyzing gene sets and building topologically-aware analysis approaches.

The MetExplore web app is based on a Java library called met4j [84]. Both MetworkPy and met4j can be used to construct and analyze metabolic networks in their respective software ecosystems. MetworkPy includes additional topological analysis features not available in met4j. These include density, neighborhood enrichment, and methods for analyzing multiple target sets (e.g. clustering, fuzzy overlap, and target set networks). In addition, MetworkPy includes flux sampling analysis tools not available in met4j.

An approach similar to network neighborhoods MetworkPy implements is used by [43] in identifying “reporter metabolites.” MetworkPy includes methods for creating neighborhood-based analyses that generalize neighborhood definitions by allowing user-selected radii and usage on weighted graphs.

#### Flux sampling analysis

There are several packages available for performing flux sampling using variations of the hit-and-run algorithm, including: COBRA [85], COBRApy [86], LooplessFluxSampler [87] and dingo [88]. Additionally, code is available for the corner sampling performed in [59]. MetworkPy includes a corner-based sampler that is easier to use for alternate analyses than is currently available from [59]. Further, MetworkPy includes tools for analyzing flux samples (obtained from any sampling approach) not available in the other packages. Among studies that analyze the results of flux sampling, they have generally only compared the marginals for single reactions [17, 33, 34, 59, 61, 62, 89], including those that use the Kullback-Leibler divergence [59, 61, 62, 89]. MetworkPy’s implementation of the Kullback-Leibler and Jensen-Shannon divergence metrics uses a nearest-neighbor based approach (based on [65, 66]), which can generalize to higher dimensions. Importantly, MetworkPy’s implementation enables direct comparison of groups of reactions via FSDA.

#### Metabolite subnetwork analysis

Previous studies have defined metabolite synthesis [24, 25, 90] and consumption [90] subnetworks to investigate metabolic perturbations, primarily to extract metabolic data from differential gene expression data. The identified metabolite networks from DPA [24, 25] (called producibility plots) include all reactions required to produce a particular metabolite, while MetPaths [90] are local, including only reactions which are proximate to the metabolite. MetworkPy defines the metabolites subnetworks globally, similar to [24, 25], and generalizes the producibility plots to allow for weighting with pFBA [49] or gFBA [50]. MetworkPy also can identify reactions and their associated genes that consume a given metabolite or its products. The code used to generate the synthesis networks in [24, 25] is not available, and the code for creating the MetPath [90] metabolite subnetworks is currently only available as a MATLAB toolbox. The metabolite networks were used with differential expression data. In the studies [24, 25], the networks were used in rank product tests, while in [90], the networks were used to calculate pathway scores based on weighted averages of gene expression fold-change values. MetworkPy’s metabolite networks can be used for these types of analysis as well, or with enrichment tests, or tests with other bioinformatic pipelines that accept a customized gene set input (e.g., GSEA [53], GSVA [54], etc.).

## CONCLUSION

MetworkPy is a toolkit for genome-scale metabolic contextualization of gene sets of interest without requiring optimality assumptions. By implementing complementary topology and flux sampling divergence analyses of GSMMs, MetworkPy robustly and specifically highlights reactions, pathways, and metabolic subnetworks proximal to input gene sets. MetworkPy also includes flexible, generalizable methods for creating, deconstructing, and analyzing graph representations of GSMMs to enable customizable topology-aware analysis pipelines.

MetworkPy’s objective-independent approach makes it useful for investigating a broad range of bacterial and eukaryotic systems. Moreover, MetworkPy delivers analyses that are both descriptive (identifying reactions, pathways, neighborhoods, and metabolite subnetworks that are affected by user-defined gene sets or input molecular profiling datasets) and anticipatory (predicting genetic or chemical perturbations with desired metabolic shifts with flux divergence analysis). MetworkPy analyses complement other FBA and pathway enrichment approaches to facilitate multi-level metabolism-centric hypothesis generation for both characterization and interventional applications.

## ADDITIONAL FILES

Additional File 1 (pdf): More detailed description of methods and tools available in MetworkPy

Additional File 2 (xlsx): Data related to the analyses performed on the small simulation metabolic model

Additional File 3 (xlsx): Data related to the analysis of the transcription factors of *Mycobacterium tuberculosis*

## AVAILABILITY AND REQUIREMENTS

Project name: MetworkPy

Project home page: https://github.com/Ma-Lab-Seattle-Childrens-CGIDR/metworkpy

Operating system(s): Platform independent

Programming languages: Python

Other requirements: Python 3.10 or higher, a compatible solver (CPLEX, Gurobi, GLPK, HiGHS, OSQP)

License: MIT

Any restrictions to use by non-academics: None

## ABBREVIATIONS

ACHR: artificial centering hit-and-run
ACGK: Acetylglutamate Kinase
ATP: Adenosine triphosphate
Arg: Arginine
Asp: Aspartate
AUC-ROC: Area under the receiver operating characteristic curve
DPA: Differential Producibility Analysis
FBA: Flux Balance Analysis
FSDA: Flux Sampling Divergence Analysis
FDR: False Discovery Rate
FVA: Flux Variability Analysis
Glu: Glutamate
Gln: Glutamine
GO: Gene ontology
GSMM: Genome-scale metabolic model
GSVA: Gene Set Variability Analysis
GUI: Graphical User Interface
H2O: water
iMAT: Integrative Metabolic Analysis Tool
Mtb: Mycobacterium tuberculosis
OCBT: Ornithine Carbamyltransferase
optGP: Optimized General Parallel Sampler
pFBA: Parsimonious Flux Balance Analysis
PYR: Purine and Pyrimidine Biosynthesis
RRA: Robust Rank Aggregation
TCA: Tricarboxylic acid cycle
TF: Transcription Factor
TFOE: Mtb strains recombinantly engineered to overexpress a transcription factor

## DECLARATIONS

### Ethics approval and consent to participate

Not applicable

### Consent for publication

Not applicable

### Availability of data and materials

MetworkPy is an open source Python package, with source code available at https://github.com/Ma-Lab-Seattle-Childrens-CGIDR/metworkpy, and can be installed from the Python Package Index (PyPI) using pip. Code used for generating the results in this paper can be found at https://github.com/Ma-Lab-Seattle-Childrens-CGIDR/metworkpy_application_note. Documentation for MetworkPy is available at https://metworkpy.readthedocs.io/en/latest/. All data associated with this study are in additional files.

## Authors’ contributions

B.G. and S.M. conceived of the methods for the Python package. B.G. implemented the package. B.G. and S.M. wrote, prepared figures for, and reviewed the manuscript.

## Acknowledgements

We gratefully acknowledge Dr. Sascha Schäuble, Dr. Jason Yang, and Mr. Ethan Bustad, for helpful discussions.

