## Additional File 1 for "MetworkPy: A Python Package for Graph- and Information-theoretic Investigation of Metabolic Networks"

**Supplementary Materials**

### S0 Glossary

- ***Stoichiometric connectivity graph***: A metabolic connectivity network derived from an input genome-scale metabolic model (GSMM). There is a bipartite form of the connectivity graph that includes both reactions and metabolites, as well as reaction-specific projections (that connect reactions which operate on a shared metabolite, either as a reactant or a product) or metabolite-specific projections (that connect metabolites that participate in shared reactions); see **Figure S2**.
- ***Metabolite subnetworks***
  - ***Synthesis***: Reactions (or their associated genes) required to produce a particular metabolite in an input GSMM (**Figure S4A**).
  - ***Consuming***: Reactions (or their associated genes) with significantly reduced maximum flux following the addition of a sink reaction that consumes all the specified metabolite generated by an input GSMM (**Figure S4B**).
- ***Reaction neighborhood***: Neighborhoods of reactions within a certain shortest-path distance of a central reaction in the stoichiometric connectivity graph (See **S2.2.2**).
- ***Density***: A metric representing the proportion of genes within a given distance of a central reaction in the stoichiometric connectivity graph that are in a target set of interest (**Figure S8**).
- ***Divergence***: A statistical distance measure. MetworkPy implements estimators for Kullback-Leibler [1] and Jensen-Shannon divergence measures [2] (See **S2.4**).

Specific MetworkPy functions mentioned in the text are colored blue.

### S1 Methods Summary

The MetworkPy package is built on top of COBRApy [3] and OptLang [4]. Thus, any of the solvers available in OptLang can be utilized by MetworkPy, including the commercial CPLEX [5] and Gurobi [6] (which both offer free licenses for academic use), as well as the open source GLPK, HiGHS [7], and OSQP [8] solvers. Additionally, any GSMM that can be handled by COBRApy can be used, including those from the BiGG database [9]. Statistical tests in MetworkPy and this article use the SciPy package [10].

The following tables provide an overview of the main functions and methods available in MetworkPy. **Table S1** catalogs functions and methods geared towards creating topological representations from GSMMs, and **Table S2** catalogs functions and methods geared towards network analysis.

**Table S1.** Graph and Subgraph creation methods implemented in MetworkPy. These methods can create graph objects that can be used with downstream analysis workflows, including pre-existing ones in MetworkPy or NetworkX, or in user-designed methods. Each row represents a different function from MetworkPy. "Function" gives the function name, "Input" specifies required user input, "Options" indicate available input arguments that users can use to modify the behavior of the function, "Output" indicates the result of the function, "Description" provides a short description of what the function does. See https://metworkpy.readthedocs.io/en/latest/ for further details. NetworkX is a Python package for working with graphs [11]. NetworkX includes a "Graph" class representing undirected graphs and a "DiGraph" class representing a directed graph. COBRApy is a Python package for constraint-based metabolic modeling [3], and includes a "Model" class representing a GSMM.

| Group | Function | Input | Options | Output | Description |
|---|---|---|---|---|---|
| **Graph Creation Methods** | create_ metabolic_ network | COBRApy Model | Can set forward and reverse reaction weights or weighting method; can set directed or undirected graph; can specify nodes to remove from graph; can specify "currency" metabolites or highly connected metabolites (i.e. metabolites that appear in a large number of reactions, e.g., ATP, $H_2O$, etc.) to remove from graph (see **S2.2.1**); can set scale functions for the product and reactant edge weights | NetworkX Graph or DiGraph | Create a bipartite stoichiometric connectivity graph representation of the GSMM. Includes nodes representing both reactions and metabolites, and reactions connected to their substrates, and flexible edge weights. |
| | create_ reaction_ network | COBRApy Model | Includes options from 'create_metabolic_ network'; can optionally incorporate arbitrary functions for combining two edge weights (the edge from a reaction to a metabolite, and from that metabolite to another reaction); can incorporate arbitrary functions for combining multiple edge edges (when two reactions share multiple metabolite substrates). | NetworkX Graph or DiGraph | Create a reaction connectivity graph representation of the GSMM. Nodes represent reactions and are connected based on whether they share substrates. This is a projection of the bipartite metabolic network onto the reaction nodes. |
| | create_ metabolite_ network | COBRApy Model | Includes options from 'create_metabolic_ network'; can optional arbitrary functions for combining two edge | NetworkX Graph or DiGraph | Create a metabolite connectivity graph representation of the GSMM. Nodes represent metabolites |

| | | | | | |
|---|---|---|---|---|---|
| | | | weights (the edge from a metabolite to a reaction, and from that reaction to another metabolite); can incorporate arbitrary functions for combining multiple edge edges (when two metabolites share multiple reactions). | | and are connected based on whether they are substrates of the same reaction. |
| | create_gene_ network | COBRApy Model | Can set graph to be directed or undirected; can specify currency metabolite groups or highly connected metabolites to remove from graph; can specify nodes to remove. When connecting genes associated with two neighboring reactions, should those genes be required to be essential for the reactions in order to be connected, or should all associated genes be connected. | NetworkX Graph or DiGraph | Create a gene connectivity graph representation of the GSMM. Nodes represent genes in the GSMM, and are connected if they are associated with the same reaction, or neighboring reactions. |
| | create_target_ set_ neighborhood_ network | NetworkX Graph or DiGraph, Target sets of nodes | Can set optional edge attribute to use for edge weight, linkage method (minimum, maximum, or mean); can set graphs to be directed or undirected; can specify maximum distance for nodes to be considered neighbors; can specify method for weighting edges (see **S2.2.1**) | NetworkX Graph or DiGraph | Create a target node set neighborhood network from a Graph (or DiGraph), and a user-provided set of target node sets. The nodes in this new network represent target sets of nodes in the original network and are connected if the two target sets share nodes or have nodes within a specified max distance of each other. |
| | create_target_ set_distance_ network | NetworkX Graph or DiGraph, target sets of nodes | Can set linkage method for summarizing group distance (minimum, maximum, or mean); can set directed or undirected graph; can set optional edge attribute to use for edge weight | NetworkX Graph or DiGraph | Create a target set distance network from a Graph (or DiGraph), and a user provided set of target node sets. The nodes in this new network represent target sets of nodes in the original network and have edges between them based on the |

| | | | | | |
|---|---|---|---|---|---|
| | | | | | aggregated (minimum, mean, maximum) distance between the nodes in the two target sets in the original Graph. |
| | create_mass_flow_graph | COBRApy Model | Can set directed or undirected graph; can specify nodes to remove; can specify number of highly connected metabolites to remove; can optionally add flux weights to edges; can set function for scaling the edge weights | NetworkX Graph or DiGraph | Create a graph representing the mass flow between reactions in the GSMM (based on [12]). Nodes are reactions in the GSMM, and edges are weighted by mass flow, based either on stoichiometry or flux. |
| | create_metabolite_mass_flow_graph | COBRApy Model | Can set directed or undirected graph; can specify nodes to remove; can specify number of highly connected metabolites to remove; can optionally add flux weights to edges; can set function for scaling the edge weights | NetworkX Graph or DiGraph | Create a graph representing the mass flow between metabolites in the GSMM (modification of the method in [12], applied to metabolites instead of reactions). Nodes are metabolites in the GSMM, and edges are weighted by mass flow, based on either stoichiometry or flux. |
| | create_adjacency_matrix | COBRApy Model | Can set forward and reverse reaction weights or weighting method; can set directed or undirected graph; can specify currency metabolite groups to remove; can set scale functions for the product and reactant edge weights; can specify function for scaling the edge weights; can return object type | NumPy array, or SciPy sparse array, or Pandas DataFrame (depending on user selection) | Create an adjacency matrix of a bipartite graph (directed or undirected) representation of the GSMM. The represented graph includes nodes for both reactions and metabolites, with edges connecting reactions and substrates with flexible weights. |
| **Network Neighborhoods** | get_graph_neighborhoods | NetworkX Graph or DiGraph | Can specify desired radius of neighborhood | dict of node ids to sets of node ids | Get the neighborhoods around each node in a Graph or DiGraph |
| | get_graph_gene_neighborhoods | NetworkX Graph or DiGraph, COBRApy Model | Can specify desired radius of neighborhood; can set whether genes should be essential for reactions in the neighborhood | dict of node ids to sets of gene ids | Get the genes in neighborhoods around each node in a Graph or DiGraph |

| | | | | | |
|---|---|---|---|---|---|
| | graph_ neighborhood_ iter | NetworkX Graph or DiGraph | Can specify desired radius of neighborhood | Iterator of tuples of node id and sets of node ids | Get an iterator over neighborhoods in a Graph or DiGraph |
| | graph_gene_ neighborhood_ iter | NetworkX Graph or DiGraph, COBRApy Model | Can specify desired radius of neighborhood; can set whether genes should be essential for reactions in the network | Iterator of tuples of node id and sets of node ids | Get an iterator over genes in the neighborhoods around each node in a Graph or DiGraph |
| | get_graph_ neighborhood | NetworkX Graph or DiGraph, central node | Can specify desired radius of neighborhood | set of node ids | Get the neighborhood around a node in a Graph or DiGraph |
| | get_graph_ gene_ neighborhood | NetworkX Graph or DiGraph, COBRApy Model, central node | Can specify desired radius of neighborhood; can set whether genes should be essential for reactions in the network | set of gene ids | Get the genes in the neighborhood around a node in a Graph or DiGraph |
| | get_group_ graph_ neighborhood | NetworkX Graph or DiGraph, node group | Can specify desired radius of neighborhood | set of node ids | Get the neighborhood around a group of nodes in a Graph or DiGraph |
| | neighborhood_ map | NetworkX Graph or DiGraph, Function to map | Can specify desired radius of neighborhood; can specify whether to include central node; can set central nodes; can specify a node filter (indicating nodes to ignore); can set edge attribute to use as distance; can specify number of parallel processes to use (which can speed up computations at the cost of some additional overhead and memory usage). | dict of node id to return type of function | Map a function (in parallel) over neighborhoods in a Graph or DiGraph |
| | gene_ neighborhood_ map | NetworkX Graph or DiGraph, COBRApy Model, Function to map | Can specify desired radius of neighborhood; can specify whether genes should be essential for reactions in the neighborhood; can specify whether to include central node; can set central nodes; | dict of node id to return type of function | Map a function (in parallel) over genes in neighborhoods in a Graph or DiGraph |

| | | | | | |
|---|---|---|---|---|---|
| | | | can specify a node filter (indicating nodes to ignore); can set edge attribute to use as distance; can specify number of parallel processes to use (which can speed up computations at the cost of some additional overhead and memory usage). | | |
| | get_kpaths_ subnetwork | NetworkX Graph or DiGraph, list of nodes of interest | Can specity number of shortest paths to use when constructing the subnetwork, the radius of neighborhoods to include, and the edge attribute to use as weight when finding k-shortest paths | Subgraph of the input network including the nodes of interest, and optionally paths between them, and neighborhoods around them. | Finds a subgraph of the input network which connects the nodes of interest by including the nodes of interest and (optionally) nodes lying on the k-shortest paths between the nodes of interest, and neighborhoods around each node of interest. |
| | get_gene_ kpaths_ subnetwork | NetworkX Graph or DiGraph, list of genes of interest, COBRApy Model | Can specity number of shortest paths to use when constructing the subnetwork, the radius of neighborhoods to include, and the edge attribute to use as weight when finding k-shortest paths | Subgraph of the input network including the nodes associated with the genes of interest, and optionally paths between them the nodes, and neighborhoods around the nodes. | Finds a subgraph of the input network which connects the nodes associated with the genes of interest by including the nodes and (optionally) nodes lying on the k-shortest paths between the nodes associated with genes of interest, and neighborhoods around each node associated with a gene of interest. |
| **Clustering** | get_network_ target_set_ clustering | NetworkX Graph or DiGraph, User specified groups of nodes | Number of clusters to find, Linkage function to use (minimum, maximum, mean) | The final clusters, and information on the clusters merged at each step in the clustering | Performs agglomerative clustering of target sets of nodes in the graphs based on the pairwise distances between the nodes in each group |
| | get_network_ target_set_ linkage | NetworkX Graph or DiGraph, User specified groups of nodes | Linkage function to use (minimum, maximum, mean) | A linkage matrix describing the agglomerative clustering which can be used with SciPy to create a dendrogram of the clustering | Performs agglomerative clustering of target sets of nodes in the graph based on the pairwise distances between nodes in each group, and generates a linkage matrix from the result (see SciPy documentation for more information) |

| | | | | | |
|---|---|---|---|---|---|
| Metabolite Subnetworks | find_ metabolite_ synthesis_ network_ reactions | COBRApy Model | Can specify method to use for identifying required reactions; can set subset of metabolites to find networks for; can specify return type | Pandas DataFrame or dict of metabolite id to list of reaction ids or dict of metabolite id to dict of metabolite id to float | Finds the reactions which are required to produce a particular metabolite; can alternatively find flux distributions to produce a metabolite (via pFBA[13] or gFBA [14]) |
| | find_ metabolite_ synthesis_ network_ genes | COBRApy Model | Can specify method to use for identifying required genes; can set subset of metabolites to find the networks for; can specify return type | Pandas DataFrame or dict of metabolite id to list of gene ids or dict of metabolite id to dict of gene id to float | Finds the genes that are required to produce a particular metabolite; can alternatively relate genes to flux of their associated reactions required to produce a metabolite (via pFBA [13] or gFBA[14]) |
| | find_ metabolite_ consuming_ network_ reactions | COBRApy Model | Can set subset of metabolites to find the networks for; can set whether to add sinks for other metabolites; can return type | Pandas DataFrame or dict of metabolite id to list of reaction ids | Finds the reactions that are “downstream” of a metabolite (defined as reactions with altered activity if the entire amount of metabolite produced in the model is forced into a sink pseudo-reaction) |
| | find_ metabolite_ consuming_ network_ genes | COBRApy Model | Can set subset of metabolites to find the networks for; can set whether to add sinks for other metabolites; can return type | Pandas DataFrame or dict of metabolite ids to list of gene ids | Finds the genes associated with reactions that are “downstream” of a metabolite (defined as reactions with altered activity if the entire amount of metabolite produced in the model is forced into a sink pseudo-reaction) |
| Condition-Specific Metabolic Models | imat | COBRApy Model, reaction weights | Can set value for epsilon parameter (flux cutoff for active reactions); can set value for threshold parameter (flux cutoff for inactive reactions) | COBRApy Solution | Find results of iMAT optimization problem, returning the solution |
| | generate_ model | COBRApy Model, reaction weights | Can set value for epsilon parameter (flux cutoff for active reactions); can set value for threshold parameter (flux cutoff for inactive reactions); can set method to use for creating iMAT model | COBRApy Model | Create a condition specific model based on iMAT |

| | | | | | |
|---|---|---|---|---|---|
| | ImatIter | COBRApy Model, reaction weights | Can set value for epsilon parameter (flux cutoff for active reactions); can set value for threshold parameter (flux cutoff for inactive reactions); can specify desired outputs to return from each iMAT model | Reaction activities (active/ inactive) or Models generated based on iMAT solutions | Class for iterating through sequential iMAT solutions, using integer-cut constraints to ensure each solution is different, with option to maximize the distance between sequential iMAT solutions, or to use a randomized objective to find diverse solutions |

**Table S2.** Analysis methods and pipelines to investigate the graphs and subgraph objects generated by MetworkPy, to contextualize gene sets of interest within metabolism. Each row represents a different function from MetworkPy. “Function” gives the function name, “Input” specified required user input, “Options” indicate available options for the function, “Output” indicates the result of the function, “Description” provides a short description of what the function does. See https://metworkpy.readthedocs.io/en/latest/ for further details. The NetworkX package [11] includes a “Graph” class representing undirected graphs and a “DiGraph” class representing a directed graph. The COBRApy package [3] includes a “Model” class representing a GSMM.

| Group | Function | Input | Options | Output | Description |
|---|---|---|---|---|---|
| **Network Densities** | node_target_ density | NetworkX Graph or DiGraph, Node Targets | Can specify network neighborhood radius; can specify which nodes to find density for; can set which nodes to consider in density calculations; can specify number of processes to use | dict of node id to node target density | Find the density of targeted nodes in neighborhoods of the Graph or Digraph |
| | gene_target_ density | NetworkX Graph or DiGraph, Gene targets | Can specify network neighborhood radius; can specify which nodes to find density for; can set which nodes to consider in density calculations; can set whether to only include genes that are essential for reactions in neighborhood; can specify number of processes to use | dict of node id to gene target density | Find the density of targeted genes in neighborhoods of the Graph or Digraph |
| | find_dense_ clusters | NetworkX Graph or DiGraph, Node or gene target set | Can specify network neighborhood radius; can set the quantile threshold considered as “dense” | Pandas DataFrame of node neighborhood densities and assigned clusters | Find clusters of neighboring nodes with high node or gene target density |
| **Enrichment** | gene_target_ enrichment | NetworkX Graph or DiGraph, COBRApy Model, Gene targets | Can specify network neighborhood radius; can specify which nodes to find density for; can set which nodes to consider in density calculations; can set whether to only include genes which are essential for reactions in neighborhood; can specify which enrichment metric to return (odds-ratio or p-value), can specify number of processes to use | dict of neighborhood gene enrichment | Find the enrichment of gene targets in neighborhoods in the graph |
| | find_ metabolite_ network_ enrichment | Metabolite networks DataFrame, Target set | Can specify whether to perform false discovery rate correction | Pandas series of metabolite network enrichment values | Find the enrichment of gene targets in metabolite networks (synthesis or consuming) |

| | | | | | |
|---|---|---|---|---|---|
| **Neighborhood P-values** | combine_ neighborhood_ gene_ pvalues | NetworkX Graph or DiGraph, Mapping of Gene to p-value, COBRApy Model | Can specify mapping of genes to weights; can specify desired radius of neighborhood; can set whether genes should be essential for reactions in the neighborhood; can specify whether to include central node; can set central nodes; can specify node filter (indicating nodes to ignore); can set edge attribute to use as distance; can set number of processes to use; can set keyword arguments to pass to SciPy's combine_pvalues function | Tuple of mapping from node id to statistic returned from SciPy's combine_ pvalues [10], and mapping from node id to p-value returned from SciPy's combine_ pvalues | Combine p-values of genes in neighborhoods in a Graph or DiGraph |
| | combine_ neighborhood_ pvalues | NetworkX Graph or DiGraph, Mapping of Node to p-value, | Can specify mapping of nodes to weights; can specify desired radius of neighborhood; can specify whether to include central node; can set central nodes; can specify node filter (indicating nodes to ignore); can set edge attribute to use as distance; can set number of processes to use; can set keyword arguments to pass to SciPy's combine_pvalues function | Tuple of mapping from node id to statistic returned from SciPy's combine_ pvalues [10], and mapping from node id to p-value returned from SciPy's combine_ pvalues | Combine p-values of nodes in neighborhoods in a Graph or DiGraph |
| **Divergence** | kl_divergence | Arrays representing samples from two distributions | Can set the number of neighbors to use for estimating divergence; can specify whether the distributions are discrete; can specify which distance metric to use; can set whether to use permutation tests to calculate a p-value | Kullback-Leibler divergence between the two distributions | Finds the Kullback-Leibler divergence between two distributions represented by sample arrays |
| | js_divergence | Arrays representing samples from two distributions | See 'kl_divergence' | Jensen-Shannon divergence between the two distributions | Finds the Jensen-Shannon divergence between two distributions represented by sample arrays |
| | calculate_ divergence_ grouped | Two Pandas DataFrames representing samples from distributions | Can specify groups of columns to calculate the divergence between the two DataFrames for; can specify the type of divergence to calculate; can specify the number of parallel processes to use; can set the options | Pandas Series of divergence for the groups, or a tuple of a Pandas Series of | Finds the divergence (either Kullback-Leibler or Jensen-Shannon) between groups of columns in the two DataFrames; |

| | | | | | |
|---|---|---|---|---|---|
| | | | available for the 'kl_divergence' and 'js_divergence' functions | Divergence and a Pandas Series of p-values calculated using permutations tests | optionally finds p-values using permutation testing |
| | ko_divergence | COBRApy Model | Can specify the specific subnetworks to find the divergence for, can set the subset of genes to find the knockout divergence for, can specify the number of samples to use for calculating the divergence, can set the options available for the 'calculate_ divergence_ grouped' function | Pandas DataFrame of the divergence caused by each gene knockout, of tuple of DataFrames of the divergence and the p-values calculated via permutation testing | Finds the divergence caused by single-gene knockouts in the COBRApy Model for all the user-defined target subnetworks |
| **Centrality** | closeness_ centrality_ subset | NetworkX Graph or DiGraph, target nodes-of-interest | Can specify which nodes to find the centrality for; can set edge attribute to use for distance; can specify whether to scale metric by fraction of nodes reachable | dict of node ids to subset closeness | Finds the closeness of nodes with respect to a subset of node targets |
| | betweenness_ centrality_ subset | NetworkX Graph or DiGraph, target nodes | Can specify whether to normalize the betweenness by the number of possible paths; can set the edge attribute to use for distance | dict of node ids to subset betweenness | Finds the betweenness of nodes with respect to a targeted node subset. This function is a slightly modified version of the NetworkX function of the same name (this version has a different normalization). |
| | betweenness_ centrality_ bipartite_ subset | NetworkX Graph or DiGraph, Node partition, target nodes | Can specify whether to normalize by the number of possible paths | dict of node ids to subset betweenness (accounting for the bipartite structure) | Finds the betweenness of nodes with respect to a targeted node subset, accounting for multiple parallel paths through shared neighbors |

| | | | | | |
|---|---|---|---|---|---|
| | | | | | in one of the bipartite node sets |
| Sampling | corner_ sampling | COBRApy Model | Can specify number of samples to use; can set which reactions to use for randomized objectives; can specify whether to scale with fva; can specify number of parallel processes to use; can set seed for random number generator | Pandas DataFrame of samples | Uses the corner sampling approach of [15] to sample from GSMMs. |
| | imat_sampling | COBRApy Model, reaction weights | can set all options from 'corner_sampling' function; can set iMAT's epsilon parameter (flux cutoff for active reactions); can set iMAT's threshold parameter (flux cutoff for inactive reactions); can set fraction of optimum value constrained for the objective function | Pandas DataFrame of samples | Adds iMAT constraints to the model, and enforces the iMAT objective to be at least a user-defined fraction of the optimum value, and then uses corner sampling to sample from the model. |
| | imat_iter_ flux_sample | COBRApy Model, reaction weights | Can specify number of samples; can specify the type of sampler to use; can set iMAT's epsilon parameter (flux cutoff for active reactions), can set iMAT's threshold parameter (flux cutoff for inactive reactions), can set method to use for iterating through iMAT models | Pandas DataFrame of Samples | Iterates through sequential iMAT models (i.e. models that achieve the same iMAT objective value but with different reactions active/inactive) based on approaches in [16], and an additional approach based on corner-sampling [15] generating flux samples from each model. |

**S2 Detailed Methods**

The following section provides a summary and more detailed explanations for some of the key methods in MetworkPy, along with results from a small simulation model (described in **S2.1**) for illustrative purposes. The methods described include:

- **S2.2**: Construction of metabolic connectivity graphs and sub-graphs
- **S2.3**: Topological analysis of the metabolic graphs and sub-graphs
- **S2.4**: Analysis of flux distributions with Flux Sampling Divergence Analysis (FSDA)
- **S2.5**: Generation of condition-specific metabolic models with MetworkPy's iMAT implementation

**S2.1 Simulation Model**

To demonstrate the different methods in MetworkPy, we have defined a simple metabolic model consisting of 23 internal reactions, 24 metabolites, and 24 genes (visualized in Error! Reference source not found. as an Escher map [17]). This model includes metabolites labelled A through X (with extracellular versions of A-G, N, R, U, V, W, and X), and 7 subsystems (labelled S1-7). The metabolites flow through reactions labelled by R__<reactants>__<products>, which have associated genes which are labelled g001-g023. **Table S3** summarizes the reactions in the simulation model, including the stoichiometries and the gene-protein-reaction (GPR) associations. **Figure S1** shows a graphical visualization simulation model.

Each reaction is associated with a GPR rule, which indicates which genes are required for the reaction to be functional. These GPR rules relate reaction functionality to associated genes. GPR rules can contain:

- No genes for reactions that happen spontaneously, or do not have known gene associations
- Single genes for reactions that are the result of the action of a single gene product
- Multiple genes connected by (possibly nested) Boolean operations (AND, OR):
    - AND indicates that multiple genes are required for the reaction to function, such as is the case with a multiple-enzyme complex
    - OR indicates that multiple genes can allow for the reaction to proceed, such as in the case of isozymes

For example, reaction R_C_H__I in the simulation model (third row in **Table S3**) requires 'g002 or g003', indicating that either gene g002 or gene g003 must be active for the reaction to proceed, wheras reaction R_F_M__N (eighth row in **Table S3**) requires 'g006 and g007,' indicating that the gene products of both g006 and g007 are required for the reaction to proceed.

**Table S3.** Summary of the reactions defined in the simulation model. The first column is the identifier of the reaction in the model. The second column (Equation) represents the stoichiometric equation of the reaction. The third column (Gene-Protein-Reaction Rule) describes the genes associated with the reaction. The fourth column indicates the subsystem of the reaction. This table does not include the exchange pseudo-reactions (these reactions are included in the 'Model Information' sheet of **Additional File 2**).

| | **Reaction ID** | **Equation** | **Gene-Protein-Reaction Rule** | **Subsystem** |
|---|---|---|---|---|
| 1 | biomass | U_C + W_C + X_C --> | | Biomass |
| 2 | R_A_B__G_H | A_C + B_C <=> G_C + H_C | g001 | S1 |
| 3 | R_C_H__I | C_C + H_C <=> I_C | g002 or g003 | S2 |
| 4 | R_C_D__J | C_C + D_C <=> J_C | g004 | S2 |
| 5 | R_I__P | I_C <=> P_C | g008 | S2 |
| 6 | R_J__Q | J_C <=> Q_C | g010 | S2 |
| 7 | R_E__M | E_C <=> M_C | g005 | S3 |
| 8 | R_F_M__N | F_C + M_C --> N_C | g006 and g007 | S3 |
| 9 | R_N__U | N_C --> U_C | g011 and g013 | S3 |
| 10 | R_N__T | N_C <=> T_C | g011 or g012 | S3 |
| 11 | N_exp | N_C --> N_E | | S3 |
| 12 | R_G_K__L | G_C + K_C <=> L_C | g008 | S4 |
| 13 | R_H__K | H_C <=> K_C | g009 | S4 |
| 14 | R_L__W | L_C --> W_C | g014 and g020 | S4 |
| 15 | R_O_P__R | O_C + P_C --> R_C | g015 and g020 | S4 |
| 16 | R_K__O | K_C --> O_C | g024 | S4 |
| 17 | W_exp | W_C --> W_E | g017 | S5 |
| 18 | R_exp | R_C --> R_E | g018 | S5 |
| 19 | R_P_Q__S | P_C + Q_C --> S_C | g016 | S6 |
| 20 | R_S_T__V_X | S_C + T_C --> V_C + X_C | g019 | S6 |
| 21 | X_exp | X_C --> X_E | g021 | S7 |
| 22 | V_exp | V_C --> V_E | g022 | S7 |
| 23 | U_exp | U_C --> U_E | g023 | S7 |

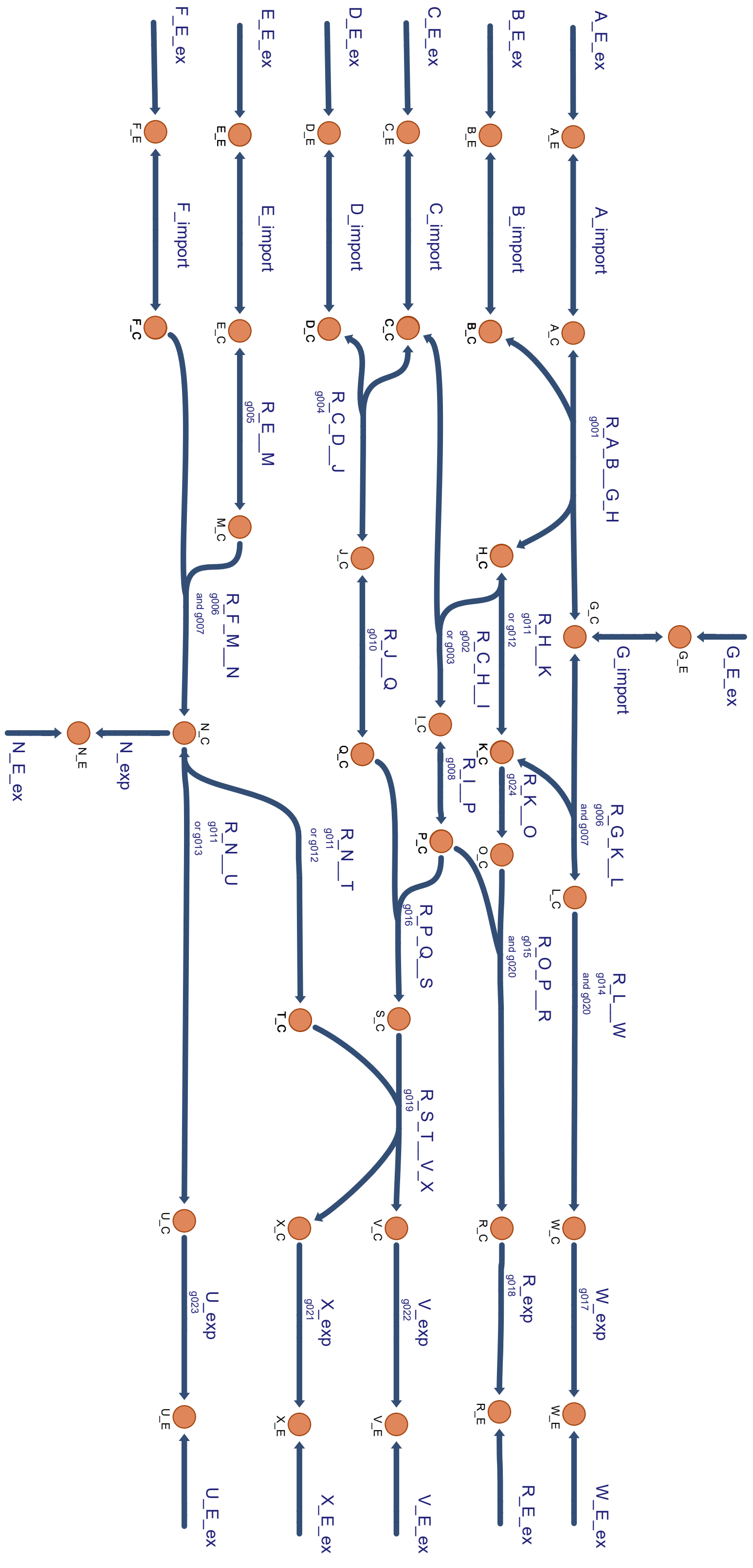


**Figure S1** A graphical representation of the Simulation Model. This visualization was generated using the Escher web tool [17]. Each edge represents a reaction in the network and is labelled with both the reaction name and the associated GPR rule. The nodes represent the metabolites in the network and are labelled with the metabolite that they represent.

### S2.2 Metabolic Connectivity Graphs

#### S2.2.1 Stoichiometric Connectivity Graphs

MetworkPy's approach to constructing a graph representation of a GSMM takes place in two stages (with the second being optional). The first step creates a bipartite graph, which can be accomplished using the create_metabolic_network function, with nodes representing both reactions and metabolites (see **Figure S2A**). This ***stoichiometric connectivity*** bipartite graph can be either directed or undirected, and either weighted or unweighted. For an undirected graph, the edges in this network represent that a reaction either consumes or produces a metabolite. For a directed graph, edges from a metabolite to a reaction indicate that the reaction consumes that, and an edge from a reaction to a metabolite indicates that the reaction produces that metabolite.

The stoichiometric connectivity bipartite graph can also be viewed as a representation of a hypergraph where the metabolites are nodes and reactions are hyperedges (i.e., edges that contain more than two nodes, in this case the metabolite substrates of the reaction). Bipartite graphs are a common method of representing hypergraphs, and all hypergraphs can be reversibly mapped to a bipartite representation [18]. The representation of a hypergraph as a bipartite graph retains the topological information from the initial hypergraph [18]. Our choice to represent GSMMs as a bipartite graph has several advantages. While packages for analyzing hypergraphs exist (e.g. HyperNetX [19] and Hypergraphx [20] in Python), the analysis tools for working with bipartite graphs are more mature and extensive. Moreover, bipartite graphs offer the ability to include different weights between a reaction and its metabolites (in a hypergraph, each reaction could only have a single weight). It can be useful to weight the edges by the metabolite's stoichiometric coefficients or by other stoichiometrically-aware weighting strategies.

MetworkPy enables flexible weighting of the edges in the stoichiometric connectivity bipartite graph. Each reaction is allowed a forward and a reverse weight, either calculated by a MetworkPy method (described below), or defined by the user (e.g., calculated from another bioinformatic tool). The forward weight is used when weighting the edges from the reaction to its products and the edges from its reactants to the reaction. The reverse weight is used for weighting the edges from the reaction to its reactants, and the edges from the products to the reaction. These weights can also optionally be multiplied by the stoichiometric coefficient of the metabolite in the reaction's equation.

For example, consider a reaction: A+3B → 2C, with a forward weight of 2.5 and a reverse weight of 5. Multiplying the weights by the stoichiometric coefficients would yield weighted edges from the reaction node to metabolite nodes A, B and C of 5.0, 15.0, and 5.0 respectively. Similarly, we could define weighted edges from A, B, and C to the reaction node as 2.5, 7.5, and 10.0.

MetworkPy has several pre-built options for weighting which can be passed to the 'weight' parameter of the "Graph Creation Methods" listed in **Table S1**:

- "stoichiometry": Edge weights are the stoichiometric coefficient of the metabolite in the reaction's equation
- "pfba": Edge weights are based on the fluxes found from parsimonious Flux Balance Analysis (pFBA)[13]. Forward weights represent the positive fluxes, and reverse weights represent the absolute values of the negative fluxes.
- "gfba": Edge weights are based on the fluxes found from geometric Flux Balance Analysis (gFBA)[14], in the same manner as for pFBA.
- "fva": Edge weights are based on the minimum and maximum fluxes for each reaction found with flux variability analysis (FVA) [21]. Forward weights are the positive maximal fluxes, and reverse reactions are the absolute value of the negative minimal fluxes.

For undirected graphs, MetworkPy specifies the weight for the edge between a reaction and a metabolite (or between a metabolite and a reaction) as the maximum of the weight for the two directed edges between the two nodes.

Currency and carrier metabolites can distort the topology of the graph. These metabolites are highly connected, but the connections are not necessarily informative about functional connections within metabolism. To address this, MetworkPy offers several strategies to either prune metabolites or metabolite reaction connections in order to avoid this distortion.

- Remove the top *n* most connected metabolites. This method finds the metabolites that are involved in the most reactions, and removes the top *n*, where *n* is user specified. This can be selected by passing *n* to the 'remove_top_metabolites' parameter of the Graph Creation Methods listed in **Table S1**.
- Remove metabolites involved in more than a certain proportion of reactions. This method finds the proportion of reactions that a particular metabolite is involved in, and if that proportion is above a user specified threshold, the metabolite is removed from the network. This can be selected by passing a float between 0 and 1 to the 'remove_top_metabolites' parameter of the Graph Creation Methods listed in **Table S1**.
- Remove balanced groups of metabolites from reactions (inspired by [22]). This approach takes a user-defined list of currency metabolite groups, which define the forms of the currency metabolites on either side of a reaction, and then for all reactions where these currency metabolites are found balanced, and they are not the only metabolites involved in the reaction, they are removed. For example, take the currency metabolite group (ATP, (ADP, Pi)) where ATP is adenosine triphosphate, ADP is adenosine diphosphate and Pi is inorganic phosphate. Reactions with ATP on one side; and ADP and Pi on the other would have the ATP, ADP, and Pi removed, unless the reaction only involved these metabolites. This can be selected by passing these metabolite groups to the 'currency_metabolites' parameter of the Graph Creation Methods listed in **Table S1.**

To construct a reaction (or metabolite) stoichiometric connectivity graph, the full metabolic bipartite graph is projected onto the reaction (or metabolite) nodes only, either internally during create_reaction_network (create_metabolite_network), or manually through bipartite_project. These can also be viewed as the line-graphs of the metabolic hypergraph and its dual [18]. See Error! Reference source not found.**B** for an example of the projection onto the reactions only (the adjacency matrix can be found in sheet 'Reaction SCN' of **Additional File 2**). Error! Reference source not found.**C** shows an example of the projection onto metabolites only (the adjacency matrix can be found in sheet 'Metabolite SCN' of **Additional File 2**).

The network projection can be either directed or undirected. Note that a directed stoichiometric connectivity bipartite graph can be projected to an undirected reaction (or metabolite) network, but an undirected stoichiometric connectivity bipartite graph cannot be projected onto a directed reaction (or metabolite) network. In the undirected case, reaction nodes are connected if they both have edges to a shared metabolite node. Similarly, metabolite nodes connect if they both have edges to a shared reaction node. In the directed case, the edges to the common node must be in opposite directions (i.e. if one edge is towards the shared metabolite node, and the other edge is away from the shared node, then the two reaction nodes will be connected). If both edges are oriented towards or away from the shared metabolite node, then the reaction nodes will not be connected.

The edges in the projected network can be weighted by the number of neighbors in the other node group that they share (e.g., a metabolite network edge can be weighted based on the number of reactions that share the metabolites as substrates), or by supplying a function which can combine the two weights of the edges to project into a single weight (e.g., minimum, maximum, mean, or sum, defaulting to minimum).

Additionally, other functions can be provided to combine the weights of multiple edges (caused by multiple shared neighbors), defaulting to maximum.

MetworkPy can create the bipartite stoichiometric connectivity network as a NetworkX [11] Graph or DiGraph through the create_metabolic_network function, but MetworkPy can also return the adjacency matrix of the network in a variety of forms via create_adjacency_matrix, including: a NumPy ndarray [23], a Pandas DataFrame [24, 25], or a SciPy sparse array [10]. The NetworkX Graph and DiGraph objects allow the user to apply NetworkX's analysis methods and for interfacing with the many Python libraries that accept NetworkX Graphs and DiGraphs. The SciPy sparse arrays can be used to interface with SciPy's sparse graph analysis functions, or with scikit-network [26].

This stoichiometric connectivity network construction and projection can be performed manually for maximum flexibility, by combining create_metabolic_network with bipartite_project, but MetworkPy also provides several wrapper functions to directly generate a reaction or metabolite stoichiometric connectivity network directly from an input COBRApy model: create_reaction_network, create_metabolite_network, and create_gene_network. The gene network is based on the reaction connectivity network and connects genes associated with the same reaction, or neighboring reactions.

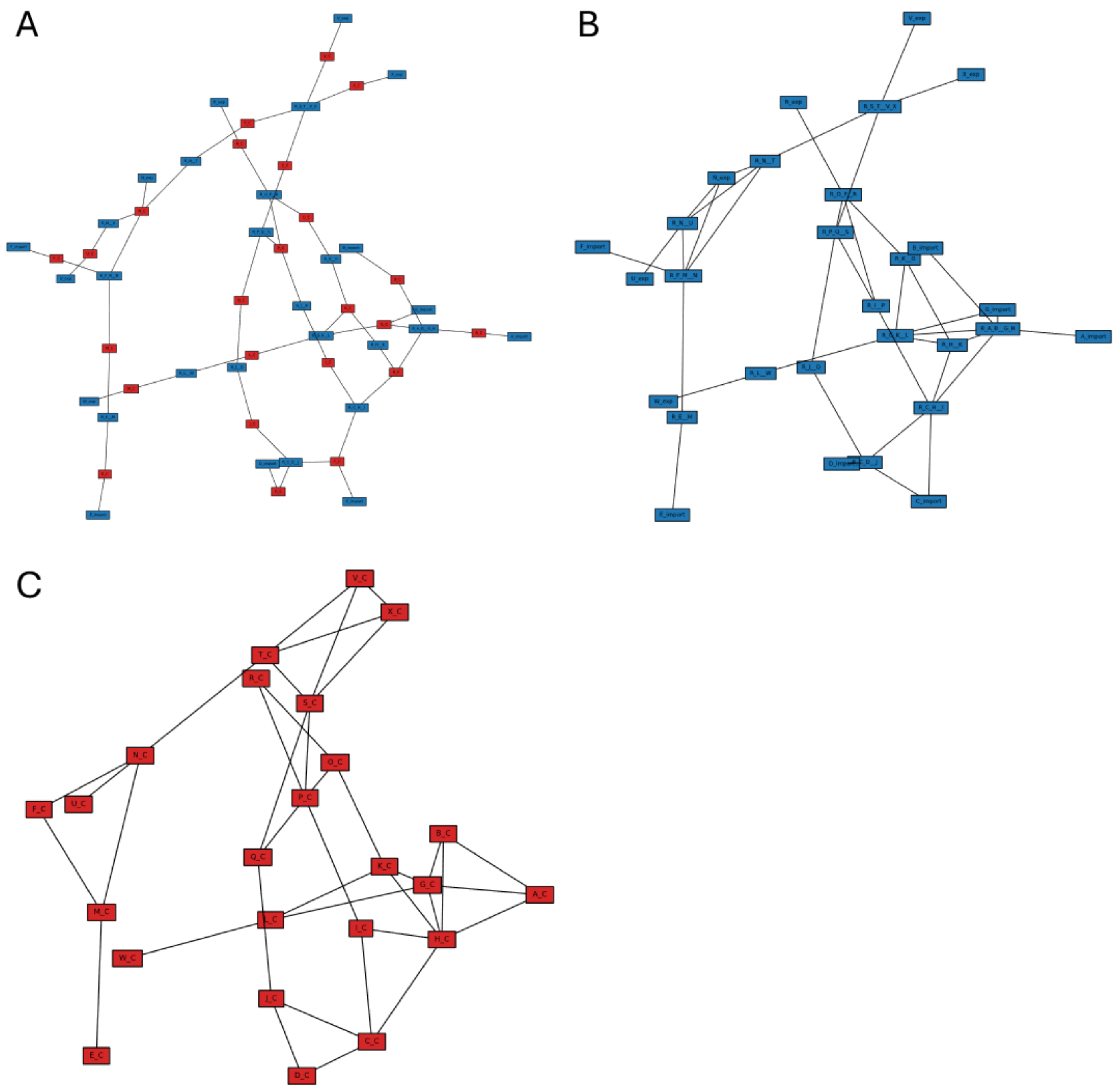


**Figure S2** Stoichiometric Connectivity Graph representations of the Simulation Model, with blue nodes representing reactions, and red nodes representing metabolites (visualized using iplotx [27]). **(A)** Shows the full metabolic network, with nodes representing reactions and metabolites. Reactions are connected to their substrate metabolites. **(B)** The bipartite metabolic network projected onto the reaction nodes only, with reactions connected if they share a substrate. **(C)** The bipartite metabolite network projected onto the metabolites, with metabolites connected if they are the substrates of the same reaction. The layout is calculated using NetworkX's 'spring_layout' function *[11]*.

In addition to the stoichiometric connectivity networks described above, MetworkPy also includes methods for constructing ***target set connectivity networks***. These networks represent the connectivity of target sets of nodes in a network. In the target set connectivity networks, each node represents a "target set" of nodes-of-interest from the original network. They are connected based on the connections between the target sets in the original network. The first type is the **target set distance network**, created with create_target_set_distance_network, which is a fully connected graph with the edge weights representing the distances between the target sets in the original network. The distance between two target sets is based on the pairwise distance between nodes in the original network, aggregated via a linkage function (e.g., minimum, maximum, and mean, which can be specified through the 'linkage' parameter). An example of this type of target set connectivity network is shown in **Figure S3A**, based on the metabolite connectivity network of the simulation model (**Figure S2C**) and target sets 'G1': {'G_C', 'I_C, 'K_C'}, 'G2': {'O_C', 'T_C'}, 'G3': {'N_C', 'E_C'}, 'G4': {'M_C', 'S_C'}, using the minimum as the linkage function.

The second type is the **target set neighborhood network**, created with create_target_set_neighborhood_network, where target sets are connected if they are neighboring in the original network. To find if two target sets are neighboring, the neighborhood of each is found (that is all the nodes within a certain maximum distance of a node in the target set). If either target set's neighborhood contains nodes of the other target set, then the two target sets are connected. The edges can also be weighted based on how many nodes from each target set lie in the neighborhood of the other; this weighting can be calculated by one of the following options selectable via the 'weighted' parameter: count (the number of nodes from the other target set that are in the neighborhood), proportion (the count divided by the size of the neighborhood), or enrichment (where a Fisher's exact test is used to calculate the enrichment of nodes in the other target set within the neighborhood). An example of a target set neighborhood network is shown in **Figure S3B**, based on the metabolite connectivity network of the simulation model (**Figure S2C**) and target sets 'G1': {'G_C', 'I_C, 'K_C'}, 'G2': {'O_C', 'T_C'}, 'G3': {'N_C', 'E_C'}, 'G4': {'M_C', 'S_C'}, with a max distance of 1 used to define the neighborhoods.

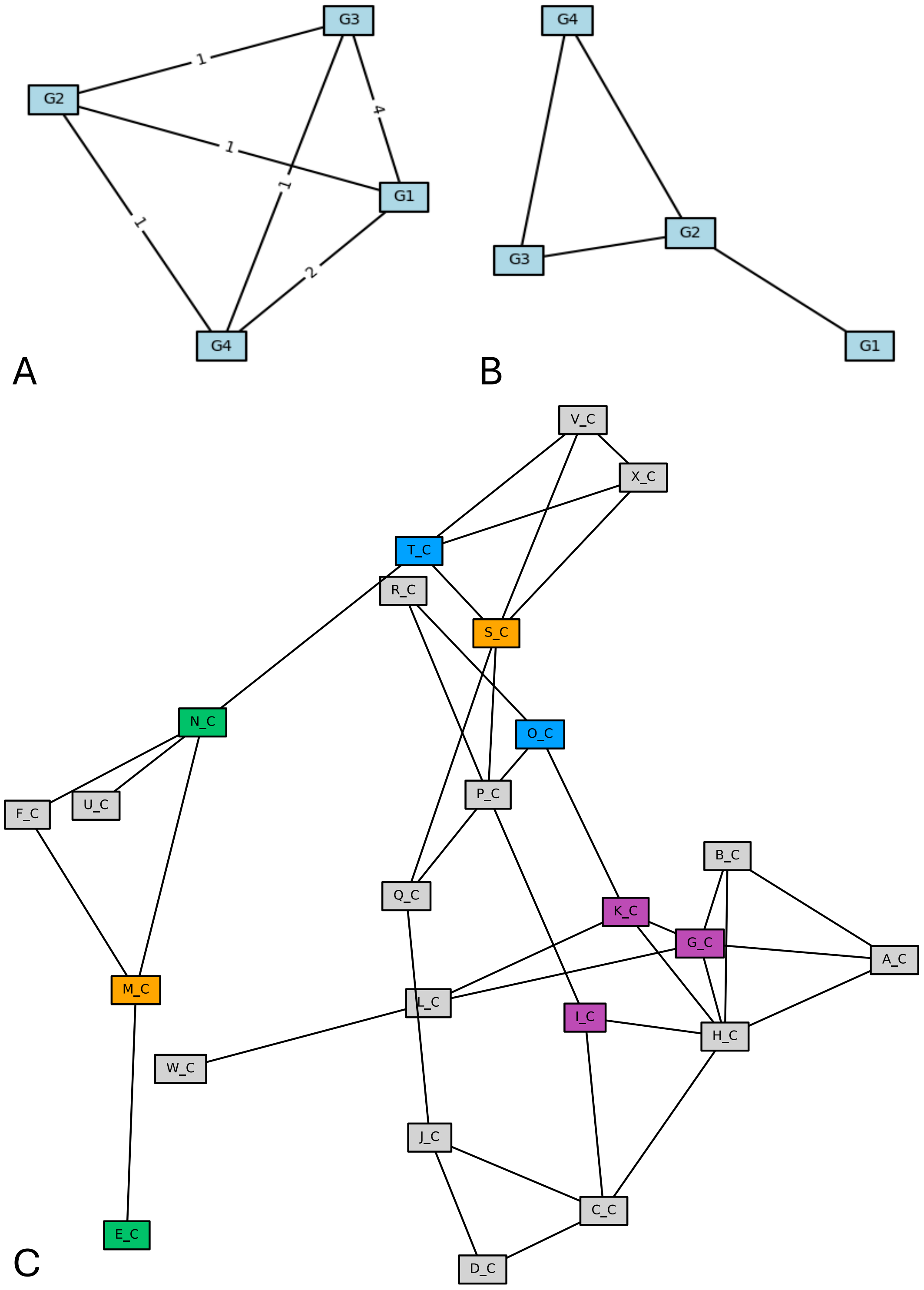


**Figure S*3*** Target set connectivity networks for metabolites in the simulation model (visualized using iplotx [27]). The target sets are sets of nodes from the metabolite connectivity network (shown in **C**), specifically 'G1': {'G_C', 'I_C, 'K_C'} colored purple, 'G2': {'O_C', 'T_C'} colored blue, 'G3': {'N_C', 'E_C'} colored green, 'G4': {'M_C', 'S_C'} colored yellow. **(A)** Shows the target set distance network created based on the distances between nodes in the two target sets, specifically using the minimum distance between a node from each target set. **(B)** Shows the target set neighborhood network created based on whether nodes from the two target sets were within a maximum distance (here 1) of each other in the original network.

#### S2.2.2 Reaction Neighborhoods

MetworkPy defines subnetworks in the stochiometric connectivity graph via ***reaction neighborhoods***. For each reaction in the graph, its neighborhood is defined as the set of reactions (or associated genes) that have shortest path distances of at most 'radius' (specified by the user) to the reaction defining the neighborhood. MetworkPy includes a variety of methods for identifying these subnetworks, iterating over them, and mapping functions (in parallel) over them (see **Table S1** or MetworkPy's documentation for more information). These methods are intended to allow for easy user-defined development of custom analysis approaches making use of network neighborhoods.

The neighborhood-based methods defined in MetworkPy can handle both the full metabolic stoichiometric connectivity graph (which includes both reaction and metabolite nodes) and the projected reaction/metabolite stoichiometric connectivity graphs. The methods for interacting with the neighborhood subnetworks allow for selecting which nodes can act as neighborhood centers. This allows, for example, only analyzing neighborhoods in the bipartite stoichiometric connectivity graph that are centered on nodes associated with metabolites. The nodes which are part of a neighborhood can also be filtered, independently of the neighborhood definition. For example, neighborhoods can be filtered to only include nodes associated with reactions.

#### S2.2.3 Metabolite Subnetworks

MetworkPy defines two types of metabolite subnetworks: 1) ***synthesis networks*** (the subnetwork of reactions and/or associated genes that are required to generate a particular metabolite) created with find_metabolite_synthesis_network_reactions and find_metabolite_synthesis_network_genes, and 2) ***consuming networks*** (the subnetwork of reactions and/or associated genes that consume the metabolite or products of the metabolite); created with find_metabolite_consuming_network_reactions and find_metabolite_consuming_network_genes, the consuming network corresponds approximately to reactions that will be impacted if the metabolite in question is depleted and removed from the model.

To determine both types of metabolite subnetwork, a sink pseudo-reaction is added to the GSMM, which absorbs the metabolite and removes it from the model. For the synthesis network, the flux through this pseudo-reaction is then set as the objective function, and the maximum flux for the generation of that metabolite is then identified by optimizing the flux through this sink pseudo-reaction using flux balance analysis (FBA) [28]. The reactions (or associated genes) that are required to generate this metabolite are then identified by performing essentiality analysis, e.g. by *in silico* knocking out each individual reaction (or gene) and determining if this significantly reduces the maximum possible flux through the metabolite sink pseudo-reaction.

To identify the reactions that consume the metabolite or its products (or reactions that would be impacted by depletion of a metabolite), an additional constraint is added to the GSMM, which forces any generated flux for the metabolite to flow into the metabolite sink pseudo-reaction, essentially absorbing all of the metabolite-of-interest from the GSMM. The maximum flux (in both directions) for each reaction is then compared between this depleted model and the unconstrained reference. If the maximum flux through a reaction is significantly reduced, then this indicates that the reaction requires the metabolite-of-interest to be active, likely because the reaction is consuming the metabolite or a product generated from the metabolite.

For both synthesis and consuming networks, sinks for other metabolites in the model can optionally be added in order to avoid artifacts caused by blocked reactions. For example, in the case of synthesizing metabolite 'X_c' in the simulation model, the reaction 'V_export' could be identified as required to synthesize 'X_c' since without being able to remove the accumulation of 'V_c', reaction 'R_S_T__V_X' would become blocked due to the steady state assumptions of FBA. Similar issues could arise in the case of consuming metabolite networks. Adding sink reactions for these other metabolites allows for isolating reactions that are directly up- or down-stream of a metabolite.

Additionally, the reactions/genes that can be included in each metabolite network can be filtered using the ‘reaction_filter’ or ‘gene_filter’ parameters of the metabolite network functions. These filters indicate which reactions or genes to check for being part of the network and can be combined with the network neighborhood functions (i.e. with get_graph_neighborhoods or get_graph_gene_neighborhoods) to only include reactions within a certain distance of the metabolite to find local metabolite networks.

These metabolite networks can be directly used to identify reactions (and their catalyzing gene products) that are associated with either the biosynthesis or consumption of a metabolite-of-interest. The metabolite subnetworks can also be combined with a target gene set from an independent bioinformatic analysis (e.g. differential gene expression) to predict metabolite signatures associated with the gene set of interest. Ultimately, these metabolite networks allow for topology-driven decomposition of the GSMM into subsets that can be analyzed in many possible ways (e.g., enrichment analysis, described in **S2.3.2**; gene set variation analysis [29]; etc.), enabling metabolic contextualization of a variety of bioinformatic methods.

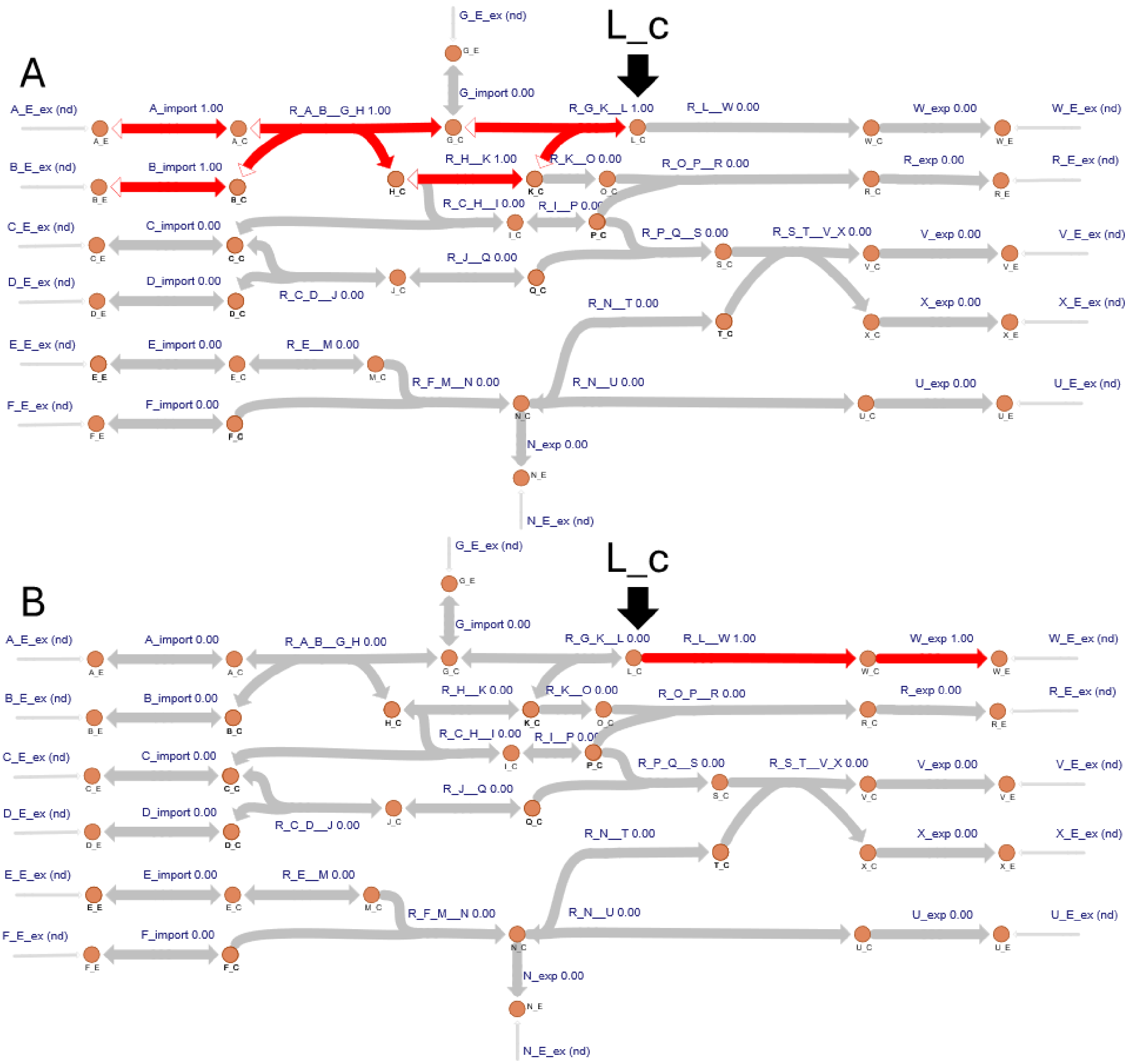


**Figure S4** Metabolite subnetworks associated with the metabolite L_c in the simulation model visualized using the Escher web tool [17]. Edges associated with reactions included in the subnetwork shown in red, and all other reactions shown in gray. The exchange pseudo-reactions are not included in either the metabolite synthesis or consuming subnetworks. **(A)** Shows the synthesis subnetwork, which includes all the reactions required to generate L_c. **(B)** Shows the consuming subnetwork, which includes all the reactions that have significantly decreased flux following the removal of all L_c generated from the model.

An example of the metabolite subnetworks in the simulation model is shown in **Figure S4**. The synthesis subnetwork shown in **Figure S4A** shows all the reactions involved in generating the metabolite L_c, from the import of precursors through several transformations to finally generate the metabolite. The fate of this metabolite is then shown in **Figure S4B**, which shows both a reaction directly consuming the metabolite, but also a reaction that acts on a product of this previous reaction. **Additional File 2** includes the metabolite synthesis and consuming networks of the simulation model in sheets ‘Metabolite Synthesis Networks’ and ‘Metabolite Consuming Networks,’ respectively.

### S2.3 Network Analysis

#### S2.3.1 Calculating Graph-theoretic Properties

Once a metabolic connectivity graph has been generated using MetworkPy, graph-theoretic analysis can be applied using NetworkX [11]. An important metric is the centrality of nodes within the metabolic connectivity graph. Centrality measures quantify the importance of nodes to the network based on their position within a network. The centrality of nodes can be evaluated with a variety of metrics, but two commonly used metrics are the "closeness centrality" and the "betweenness centrality."

The closeness centrality describes how central a node is in terms of the average shortest path distance to other nodes in the network, with higher values indicating that a node has shorter average path lengths to other nodes in the network. The closeness centrality of reactions and metabolites in the simulation model is shown in **Figure S5A** (**Additional File 2** includes the full set of the simulation model's reaction centrality values for the reactions in sheet 'SCN Reaction Centrality', and the metabolites in sheet 'SCN Metabolite Centrality'). The reactions near the core region of the metabolic connectivity graph have higher closeness centrality values, acting like a hub with potentially many different fates for the metabolites that these reactions produce. Regulation in regions like this can have significant impacts on the fates of core metabolites.

The betweenness centrality describes how central a node is in terms of the proportion of shortest paths between nodes in the graph that pass through it. The reactions with high betweenness centrality are those that connect two distinct parts of the metabolic network, acting similarly to bridges. If a reaction with a high betweenness centrality was inactive, it would sever a bridge between two distinct portions of the metabolism. The betweenness centrality of reactions and metabolites in the simulation model is shown in **Figure S5B** (**Additional File 2** includes the full set of the simulation model's centrality values for the reactions in sheet 'SCN Reaction Centrality', and the metabolites in sheet 'SCN Metabolite Centrality'). Reaction R_S_T__V_X (and neighbors) have a high betweenness centrality, since it connects the upper half of the simulation model (formed by the paths between A_E_ex, B_E_ex, C_E_ex, D_E_ex; and W_E_ex, R_E_ex, V_E_ex, X_E_ex) with the bottom half (formed by the paths between E_E_ex, F_E_ex; and U_E_ex). This means that for any of the shortest paths between the upper and lower parts of the simulation model, they must cross through R_S_T__V_X, giving it a high betweenness centrality. Regulation of the R_S_T__V_X reaction could activate/inactivate more distally connected components of the metabolic network (e.g. reactions involving the metabolite A).

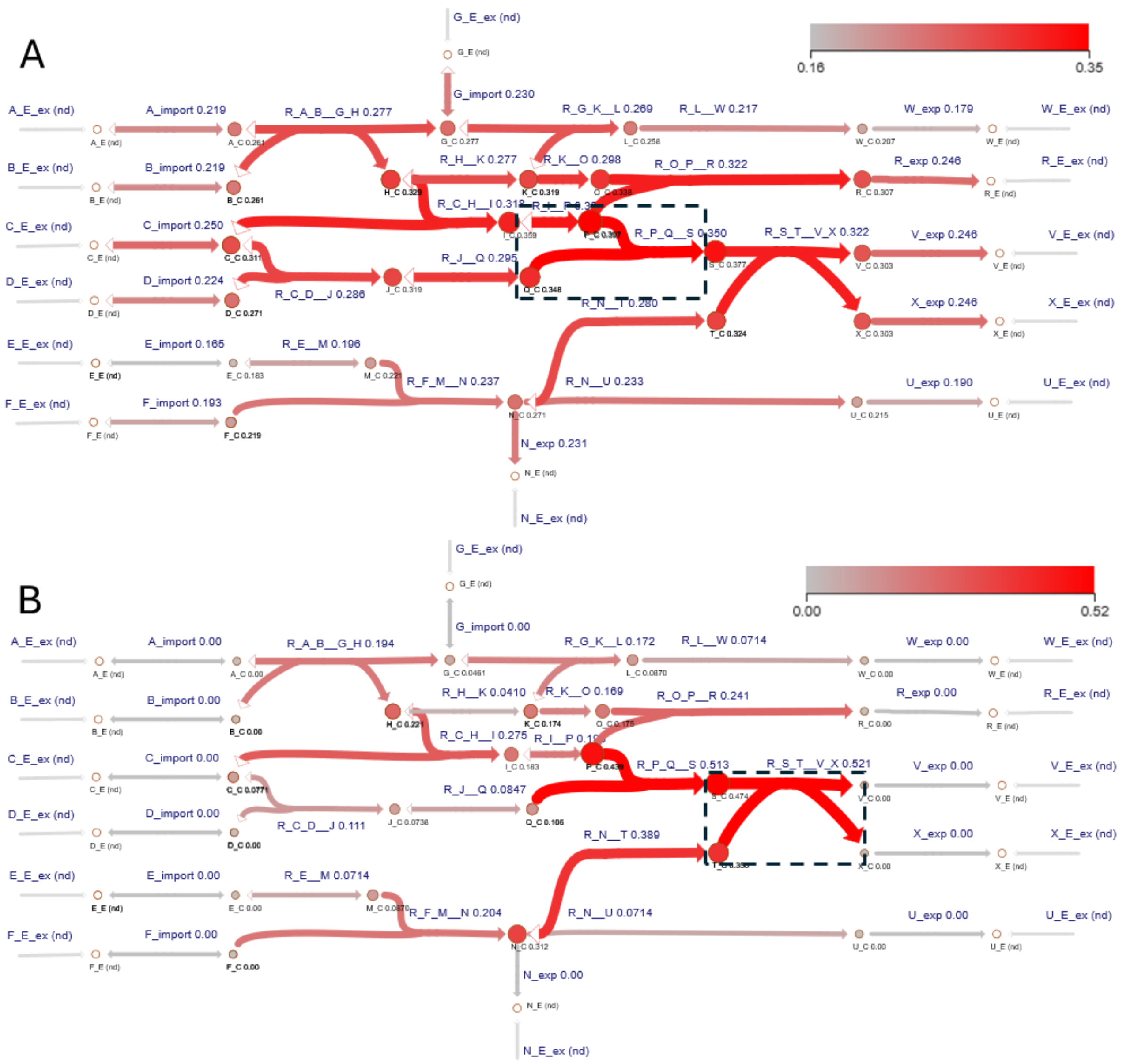


**Figure S5** Centrality measures of the reactions and metabolites in the simulation model, calculated using NetworkX [11], visualized using the Escher web tool [17]. **(A)** The closeness centrality **(B)** The betweenness centrality. The edges in the network are labelled, colored, and sized according to the centrality of the associated reaction within the reaction stoichiometric connectivity network. The nodes in the network are labelled, colored, and sized according to the centrality of the associated metabolite in the metabolite stoichiometric connectivity network. The colors of the metabolites and reactions are scaled between the minimum and maximum closeness/betweenness in the network. The most central reaction for the two centrality measures (R_P_Q__S for panel A and R_S_T__V_X for panel **B**) is marked by a dashed box.

Beyond investigating the importance of individual nodes to the full network, centrality metrics can be scoped to investigate the importance of nodes for a preselected “target” subset of nodes. The subset centrality measures can also have their non-subset metrics subtracted from them in order to identify nodes that are specifically central for the subset of interest. In the case that these nodes are regulatory targets, then the nodes that tend to fall between these targeted nodes could indicate regulation-induced metabolic rerouting.

MetworkPy includes a novel variant of the closeness centrality for a subset of nodes. This metric measures the average distance of nodes in a network to a target node set to identify nodes which are close to the node subset of interest. In the case where these nodes are reactions associated with differential gene expression; this could potentially indicate a central point of metabolic perturbation.

The subset closeness centrality for the subset of reactions {R_H__K, R_O_P__R, R_C_D__J} is shown in **Figure S6A** (**Additional File 2** includes the full set of the simulation model's subset centrality values of the reactions in sheet 'SCN Subset Centrality'). The reactions R_H_K, and R_C_H__I and the metabolites C_C, H_C, I_C, and K_C have the largest subset closeness centrality. If, for example, the targeted subset represented differential expression, this could indicate potential upstream points of regulation.

NetworkX includes a subset betweenness centrality calculation; however, NetworkX's implementation normalizes path counts to all nodes in the graph rather than to only those nodes in the subset. To adjust this calculation to fit the needs of evaluating metabolic subsets, MetworkPy includes a modification which only normalizes the subset betweenness centrality metric to the number of possible paths between nodes in the targeted subset.

In the case of the full bipartite stoichiometric connectivity network, the betweenness centrality metric can be biased by reactions that share different numbers of metabolites (since each metabolite would then be counted as a separate path between the two reactions). To address this, MetworkPy includes another variant of the betweenness centrality that focuses on the case where the subset of interest is in one of the two bipartite node subsets. MetworkPy's betweenness centrality metric counts paths through shared metabolites as one path instead of repeating it for each shared neighbor. This is equivalent to computing the betweenness on the hypergraph that the bipartite network is representing. To our knowledge, this subset betweenness centrality calculation method is novel.

The subset betweenness centrality for the subset of reactions {R_H__K, R_O_P__R, R_C_D__J} is shown in **Figure S6B** (**Additional File 2** includes the full set of the simulation model's subset centrality values of the reactions in sheet 'SCN Subset Centrality'). The subset betweenness centrality highlights both the reaction R_C_H__I, as well as the metabolites C_C and H_C. This analysis result informs the hypothesis that if the R_H__K and R_O_P__R were targeted by regulatory events, they might potentially be acting to change the reaction paths that C_C and H_C are taking, potentially rerouting them through R_C_H__I (and ultimately downstream reactions).

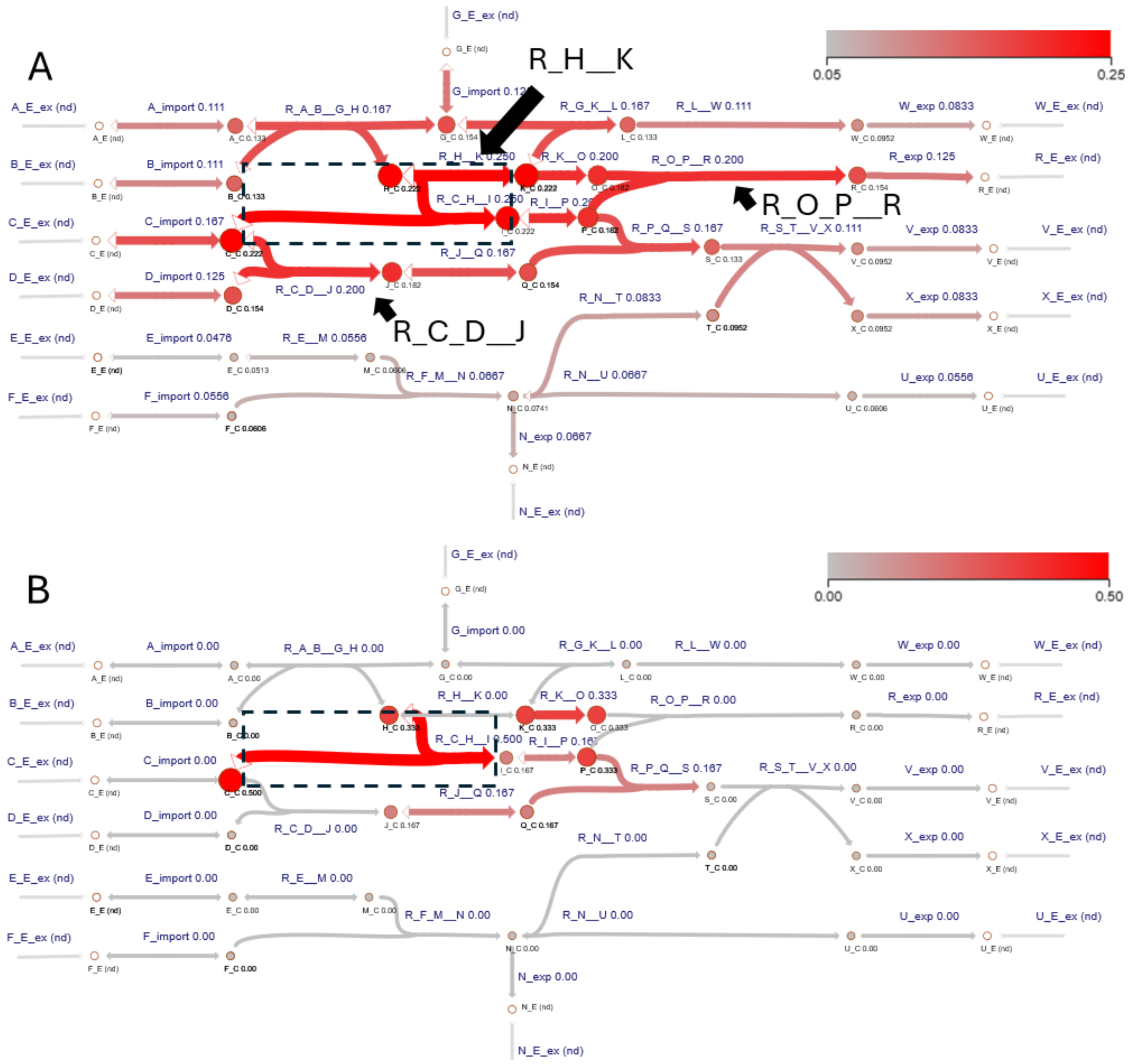


**Figure S6** Subset centrality measures of the reactions and metabolites in the simulation model, visualized using the Escher web tool *[17]* for reaction subset {R_H__K, R_C_D__J, R_O_P__R}. **(A)** The subset closeness centrality **(B)** The subset betweenness centrality. The edges in the network are labelled, colored, and sized according to the centrality of the associated reaction within the reaction stoichiometric connectivity network. The nodes in the network are labelled, colored, and sized according to the centrality of the associated metabolite in the metabolite stoichiometric connectivity network. The colors of the metabolites and reactions are scaled between the minimum and maximum closeness/betweenness in the network. The most central reaction(s) for the two centrality measures (R_C_H__I and R_H__K for panel A and R_C_H__I for panel **B**) is marked by a dashed box.

### S2.3.2 Estimating Network Enrichment

Both metabolite subnetworks and reaction neighborhoods can be evaluated for the enrichment of a target set of genes, using the find_metabolite_network_enrichment and gene_target_enrichement functions respectively. The genes associated with these subnetworks are defined by the GPR rules from the GSMM. Genes in a reaction's neighborhood are then analyzed for enrichment using a Fisher's Exact test. MetworkPy can either return the p-value of the test or the odds-ratio (based on user-input).

Enrichment of gene targets in metabolite subnetworks or reaction neighborhoods is useful in identifying metabolic overlaps between multiple gene target sets. MetworkPy's enrichment analysis can identify metabolic overlaps that do not require exact intersection between the target sets. An example of how this analysis could be useful is in the case of examining multiple gene regulators that share a common phenotype. Metabolic overlaps of their gene targets (and associated reactions) could point to a potential shared metabolic correlate of the common phenotype. Gene sets derived through different bioinformatic pipelines can also be integrated in a similar way. For example, sets of differentially expressed genes measured from multiple conditions could be overlapped to inform hypotheses about shared metabolic responses between the distinct conditions.

To perform these types of overlaps, enrichment (or density; discussed in **S2.3.3**) can be evaluated for the metabolite subnetworks or reaction neighborhoods for multiple gene sets. For each gene set, this defines a fuzzy set on the reactions, with decreasing membership for reactions that are not enriched or with low density. The results of these analyses can then be combined, for example using rank aggregation, to identify any metabolic overlaps between the multiple gene sets. MetworkPy also includes a submodule for performing this type of fuzzy overlap. The module allows users to apply different membership functions, including ones based on distances to the *k*th nearest member of the gene set, distance weighted densities, subset closeness, subset betweenness. Alternatively, a user-defined function can be used to determine the membership. The fuzzy set membership is determined using the fuzzy_reaction_set function. Once the fuzzy reaction set is defined, with each reaction being assigned a membership between 0 and 1, multiple of these fuzzy sets can be overlapped using the fuzzy_reaction_intersection function. Overlap functions include fuzzy intersection (minimum), fuzzy union (maximum), or alternate approaches based on rank aggregation (e.g., geometric mean, Robust Rank Aggregation [30], or p-value combination). See online documentation at https://metworkpy.readthedocs.io for more information.

**Figure S7** shows an example of the reaction-neighborhood enrichment in the simulation model using different values for 'radius'. As the radius increases, reactions with some enrichment in their neighborhood spread out across the network, but the significance decreases. The enrichment p-values for all the reactions in the simulation model can be found in the 'Gene Target Enrichment' sheet of **Additional File 2**.

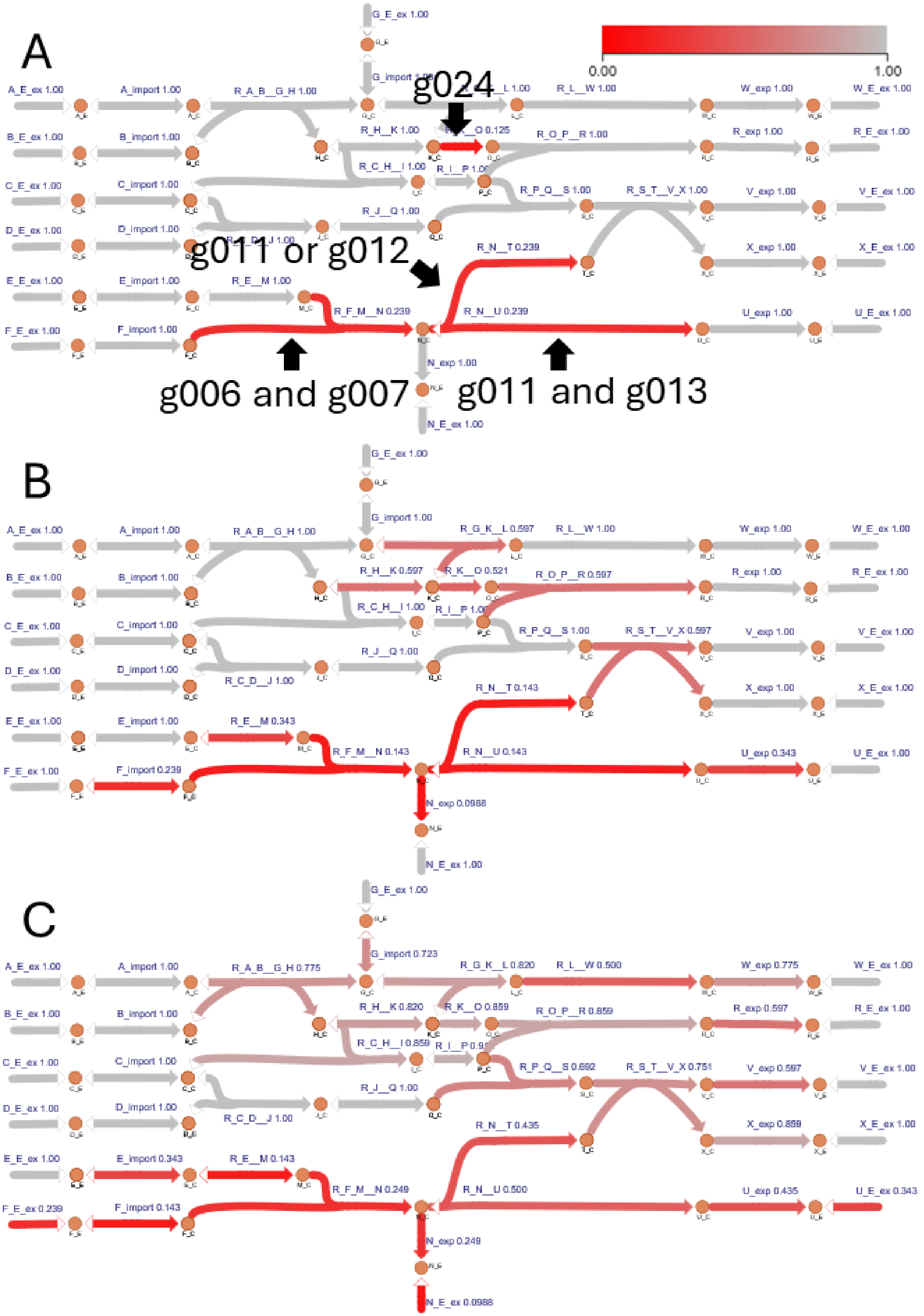


**Figure S7** Gene target enrichment p-values calculated for the target gene set {'g006', 'g011', 'g024'} on reaction neighborhoods with a specified radius of 0 **(A)**, 1 **(B)**, or 2 **(C)**. A reaction neighborhood of radius 0 includes only the reaction itself; neighborhoods of radius 1 include both the reactions and direct neighbors in the reaction stoichiometric connectivity network. The edges in the network are labelled and colored based on the p-value of enrichment for the gene targets in the neighborhoods around the associated reactions. P-values close to 0.0 are more red, and those near 1.0 are shown in grey. This network is visualized using the Escher web tool [17].

#### S2.3.3 Calculating Density

Enrichment metrics can struggle in situations where there are a large number of gene targets outside of the overlap region, since these irrelevant genes decrease the significance of enrichment in a specific region. To address this, MetworkPy includes a ***density*** metric, through the node_target_density and gene_target_density functions, which is defined as the proportion of genes (or reactions) in a subnetwork that are in the gene target set. This metric is not impacted by genes outside of the subnetwork. It is thus able to identify local increases in density without regard to the size of the total gene set. The density metric is useful in identifying overlaps in the case of a large number of genes outside the overlap region of interest.

MetworkPy offers several methods of evaluating the enrichment or density of gene targets in reaction neighborhoods:

1. Targeted Reaction Density: Gene targets are translated into reaction targets using GPR rules. The density is the proportion of reactions in a neighborhood that are targeted.
2. Target Gene Density: Gene targets are used directly, and the reactions in a neighborhood are associated with genes using GPR rules. The density is the proportion of the genes in the neighborhood that are targeted.

An example of the reaction neighborhood density for the target gene set {‘g006’, ‘g011’, ‘g024’} with different values of radius is shown in **Figure S8**. With a radius of 0, the only reactions with any density are those directly associated with the gene targets. As the radius of the neighborhood increases, the density spreads across neighboring reactions. The density values for all of the reactions in the simulation model can be found in the ‘Gene Target Density’ sheet of **Additional File 2**.

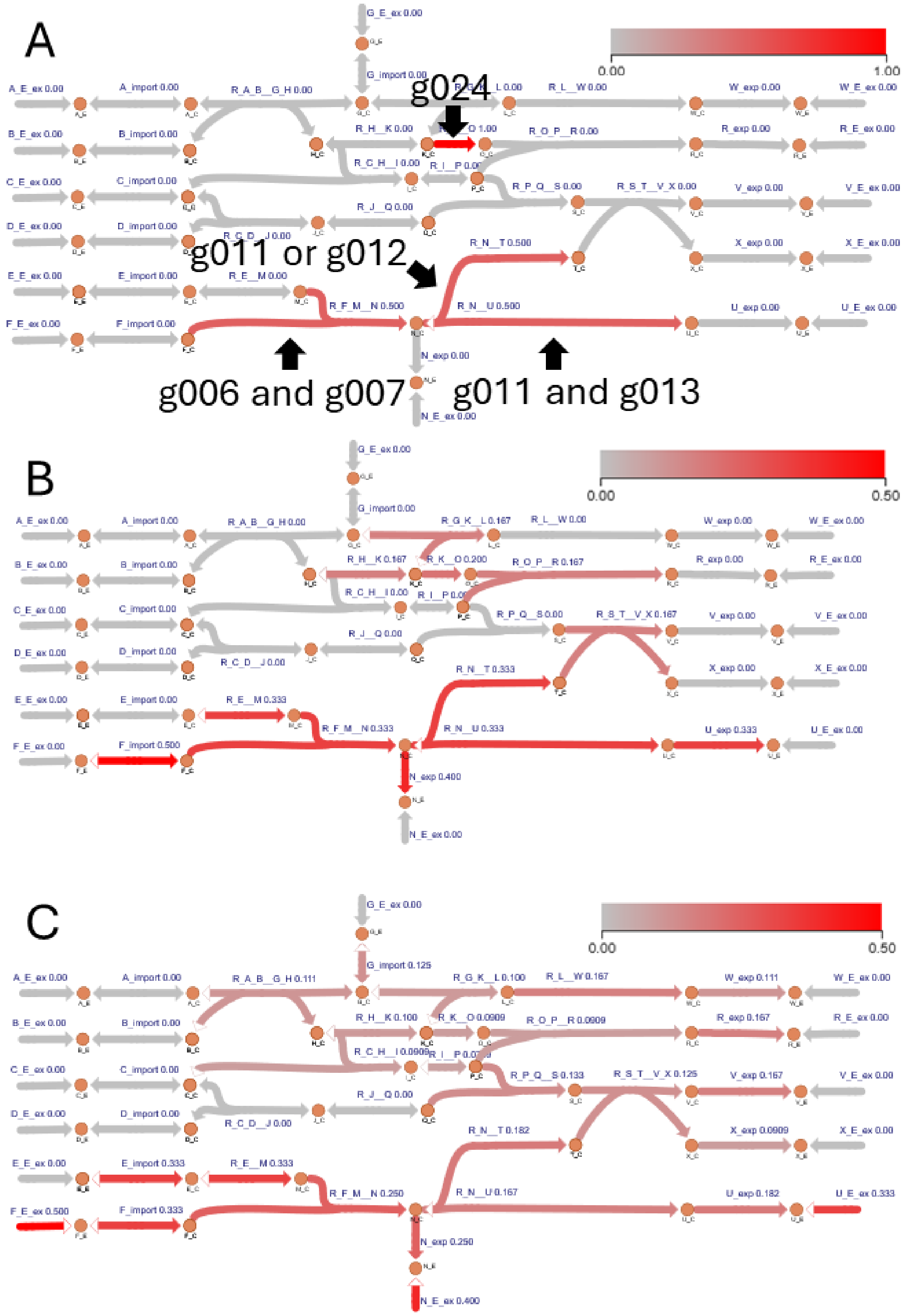


**Figure S8** Gene target densities calculated for the target gene set {'g006', 'g011', 'g024'} on reaction neighborhoods with a specified radius of 0 **(A)**, 1 **(B)**, or 2 **(C)**. A reaction neighborhood of radius 0 includes only the reaction itself; neighborhoods of radius 1 include both the reactions and direct neighbors in the reaction stoichiometric connectivity network. The edges in the network are labelled and colored based on the density of the gene targets in the neighborhoods around the associated reactions. Densities close to 1.0 are more red, and those near 0.0 are shown in grey. This network is visualized using the Escher web tool [17].

**S2.3.4 Target Set Clustering**

In addition to the fuzzy overlap described above (**S2.3.2** and **S2.3.3**), MetworkPy can also facilitate the clustering of target sets of nodes. This can assist in identifying metabolic similarities between target sets. For example, a given set of regulators could be clustered based on the proximity of their regulatory targets in the stoichiometric connectivity graph. Such analysis would posit potentially related regulators. Target node clustering can be done in several ways with MetworkPy.

The first is to calculate the aggregated distance matrix (using create_target_set_distance_adjacency_matrix) between target sets. This method finds the distance between target sets based on the pairwise distance between all nodes in each set, and then aggregating those distances (using e.g. minimum, maximum, mean). This aggregated distance matrix can then be used with any of scikit-learn's [31] clustering algorithms that allow for precomputed distance matrices (e.g. AgglomerativeClustering, DBSCAN, and HDBSCAN). However, the aggregated distance matrix approach cannot update the distances between groups based on the underlying nodes.

To address this limitation, MetworkPy also includes direct agglomerative clustering (get_network_target_set_clustering), which will recompute this target distance matrix during the agglomerative clustering process based on the underlying nodes of the target sets. This result can also be returned in the form of a SciPy linkage matrix (get_network_target_set_linkage) which can be used with SciPy's dendrogram function to generate a dendrogram of the target node sets.

**S2.3.5 Estimating Neighborhood P-values**

MetworkPy also provides a method for aggregating p-values for genes in the neighborhoods of a graph. Specifically, the function combine_neighborhood_pvalues uses SciPy's 'stats.combine_pvalues' function to aggregate p-values in neighborhoods of the metabolic graph. Several approaches for aggregation are available, including Stouffer's, Fisher's, and Pearson's methods [10]. This function offers flexible filtering of the nodes considered in the metabolic network, so it can be used to recreate (and generalize) the method of [32] for identifying reporter metabolites.

**S2.4 Flux Sampling Divergence Analysis (FSDA)**

MetworkPy facilitates the comparison of flux sampling distributions through ***divergence***. Divergence is a measure of statistical distance between two distributions, and MetworkPy includes functions to estimate two divergence measures between distributions: Kullback-Leibler (KL) and Jensen-Shannon (JS) divergence. KL divergence is a measure of the expected excess surprise when using an approximation $Q$ instead of the true distribution $P$ [1], shown in **Equation 1**. The JS divergence is a modification of the KL divergence, which is the average of the KL divergences between the distributions and their mixture [2], shown in **Equation 2**. The JS divergence has the advantage of being symmetrical and guaranteeing finite values but is more computationally expensive with less direct interpretability.

$$D_{KL}(P\|Q) = \int_{-\infty}^{\infty} p(x)\log\frac{p(x)}{q(x)}\,dx$$

**Equation 1** Kullback-Leibler divergence between distribution Q and P [1]

$$D_{JS}(P\|Q) = \frac{D_{KL}(P\|M) + D_{KL}(Q\|M)}{2} \quad where\ M = \frac{P+Q}{2}$$

**Equation 2** Jensen-Shannon divergence between distribution Q and P [2]

Exact calculation of these divergence measures would require the underlying probability distributions of $P$ and Q, so for flux samples from GSMMs, the divergence must be estimated. To perform this estimation,

MetworkPy provides implementations, in the kl_divergence and js_divergence functions, of the k-nearest neighbors estimators of [33] for KL divergence and [34] for JS divergence.

In addition to calculating the divergence measures, MetworkPy also includes permutation testing for the divergence to evaluate significance. To estimate the significance of the divergence between two flux distributions, two flux samples of user-defined size from the distributions are repeatedly shuffled (a user-defined number of times) into two random groupings of the same size as the original samples. The divergence between these shuffled samples is used to estimate the null distribution. The p-value of the divergence of the original samples is then evaluated using the upper bound p-value of [35] (see **Equation 3**) so as to not overestimate significance, especially in the context of family-wise false discovery rate correction.

$$p_u = \frac{b+1}{m+1}$$

**Equation 3** Upper bound of p-value calculated via permutation testing. Where *m* is number of permutations performed, and *b* is the number of permutations with a more extreme value than the test-statistic [35].

MetworkPy can simulate the impact of gene knockouts (KOs) and multiple gene perturbations by evaluating the divergence of a flux sample from a reference GSMM to a flux sample from a perturbed GSMM that represents the *in silico* disruption of an individual or set of metabolic genes and associated reactions. The divergence is estimated for user-defined divergence groups (this can be single reactions, reaction subsystems, metabolite subnetworks, reaction neighborhoods, etc.). To aid with these analyses, MetworkPy includes a ko_divergence function, which performs single gene KOs for all genes (or a user-selected subset of genes) in a GSMM and evaluates the divergence between the GSMM with the gene KO and the reference GSMM (without the gene KO) for a set of user-defined reaction groups. Additionally, the more general calculate_divergence_grouped function can be used to evaluate the divergence for user-defined reaction groups between any reference GSMM – perturbed GSMM pair.

To demonstrate MetworkPy's divergence metrics, we used them to compare the flux space of the simulation model before and after the KO of the gene g010. The divergence resulting from this KO is shown in Error! Reference source not found.**9**. Knocking out the gene stops the associated R_J__Q reaction, and this reaction thus has high divergence. In addition to the direct reaction association, knocking out R_J__Q impacts the possible flux states of reactions both downstream and upstream of this intervention. Reactions that are highly divergent upon KO of a gene are those whose flux states are most altered by this genetic intervention. The KO divergence values for the genes in the simulation model can be found in the 'KO Divergence' sheet of **Additional File 2**.

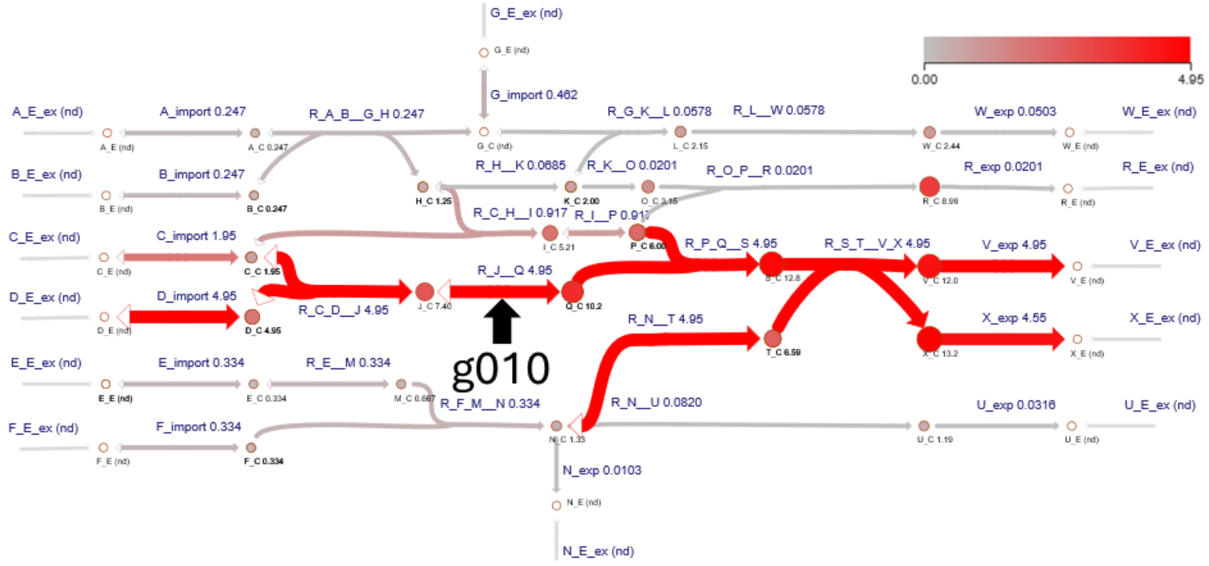


**Figure S9** Flux divergence following KO of g010 in the simulation model, visualized using the Escher web tool [17]. Shows the KL divergence between flux samples in the simulation model with g010 knocked out relative to the flux samples of the base simulation model. The edges in the graph are labelled, colored and sized based on the divergence of the associated reactions. The nodes in the network are labelled, colored and sized based on the divergence of the metabolite synthesis subnetworks. The size and colors of the metabolites and reactions are scaled between 0.0 and the maximum divergence value, with red signifying higher divergence.

#### S2.5 iMAT Implementation

The iMAT algorithm [36] is implemented in MetworkPy through the imat and generate_model functions to facilitate the analysis of condition-specific GSMMs via the other tools provided by MetworkPy. The iMAT algorithm uses gene expression data to define qualitative gene weights (-1, 0, and 1) indicating low expression levels, medium/unknown expression levels, and high expression levels, respectively. These weights can be generated using quantile cutoffs, or through differential expression analysis. Once obtained, these weights are translated into reaction weights by using the GPR rules of the GSMMs. These rules take the form of Boolean expressions between the different genes and are extended to the trinary weights by using a maximum function for ‘OR’, and a minimum function for ‘AND.’ The iMAT optimization problem uses these reaction weights as inputs, attempting to identify a flux distribution which most closely aligns with the reaction weights. It does this by defining activity thresholds: a reaction with a flux above a certain cutoff (referred to as epsilon) is considered active, and a reaction with flux below a lower cutoff (referred to as threshold) is considered inactive. The algorithm defines an optimization problem, finding the maximum number of reactions whose activity (active vs inactive) matches their weight (high expression vs low expression). MetworkPy’s implementation of iMAT allows for more general gene and reaction weights than those described above. Specifically, it treats negative weights as low-expression reactions, positive weights as high expression reactions, and zero weights as indeterminate. The binary variables in the objective function then use the absolute value of weights as coefficients. This allows using the reaction weights to express different versions of the optimization problem, for example they could be weighted by significance level of differential expression to include confidence in the iMAT problem.

The solution of the iMAT optimization problem consists of state of the binary activity variables and the flux state of the reactions in the model. MetworkPy includes several different methods of using these outputs to generate condition-specific GSMMs. These methods can be selected through the ‘method’ parameter of the generate_model function. Two of the simplest of these methods force the model to follow the activity and/or inactivity of reactions in the iMAT solution. Specifically, the ‘subset’ method knocks out all the inactive reactions, while the ‘simple’ method does this and forces the active reactions to have flux of at least epsilon.

The solution to the iMAT problem is not guaranteed to be unique; the iMAT optimization problem guarantees that the objective is a global maximum (within the solver tolerance), but there can be many possible flux distributions which result in this maximum, of which only one is found when solving. To address this, MetworkPy includes three additional methods of generating condition-specific GSMMs.

The first (called ‘imat’), adds mixed-integer constraints to the GSMM, which forces its flux distributions to be within a certain tolerance of the iMAT objective. An issue with this approach is that the model integer constraints and so cannot be sampled with the hit-and-run type samplers of COBRApy. The other two methods described below are preferable in cases where flux sampling with these methods is desired.

The second (called ‘milp’) implements the method of [37]. This approach forces each reaction in the model (or only reactions with a non-zero weight, depending on the parameter ‘constrain_all_reactions’) to be active, then inactive, checking if the iMAT objective value is greater in one case or the other. For reactions where the iMAT objective is greater for the active state, constraints are added such that they are always active, and for reactions where the iMAT reaction is greater for the inactive state, constraints are added to ensure the reaction is always inactive.

The last method (called ‘fva’) uses an approach similar to flux variability analysis, by finding the maximum and minimum fluxes through each reaction in the model (or only reactions with a non-zero weight, depending on the parameter ‘constrain_all_reactions’) while the iMAT objective function is forced to be within a user-defined tolerance of the global maximum. The minimum and maximum values of each reaction found this way are used as their new bounds to define the condition-specific GSMM.

To illustrate the use of FSDA to evaluate multiple genetic interventions, we used MetworkPy’s implementation of iMAT to generate a condition-specific metabolic model and compared it to the base simulation model using KL divergence. The genes {g014, g020, g017, and g008} were given weights of 1, the genes {g013 and g023} were given weights of -1. The solution to the iMAT optimization problem (depicting the active and inactive reactions) is shown in **Figure S10A**. After sampling from an iMAT condition-specific GSMM using MetworkPy’s ‘fva’ iMAT method, we evaluated the divergence between the iMAT model and the base simulation model for all reactions and metabolite synthesis networks. The results of this simulation are shown in **Figure S10B**. The divergence for the reactions and metabolite synthesis networks in the simulation network are reported in sheet ‘iMAT Divergence’ of **Additional File 2**.

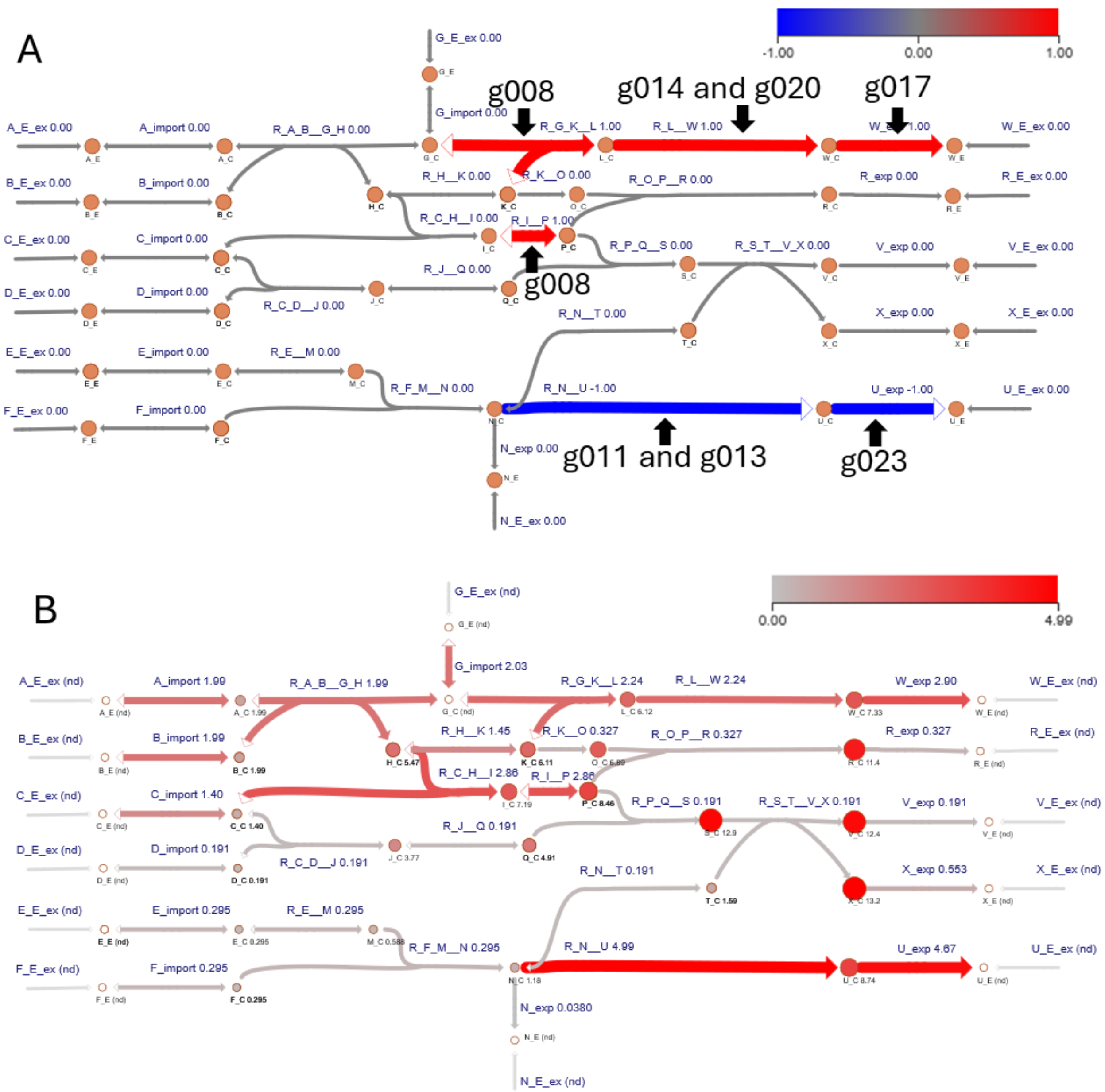


**Figure S10** iMAT [36, 37] results visualized using the Escher web tool [17]. **(A)** Shows solution to the iMAT optimization problem [36] with the reactions determined to be active by iMAT in red, and those determined to be inactive in blue, based on the binary activity variables in the iMAT optimization problem. All other reactions are colored gray. **(B)** Shows the flux divergence of the iMAT constrained model relative to the base model. The reaction color and size reflect the divergence of the associated reaction's flux distribution between the iMAT model and the base simulation model, while the metabolite color and size reflect the divergence for the associated metabolite's synthesis network. Higher divergence is shown in red.

### S3 Application to *Mycobacterium tuberculosis*

To demonstrate the utility of MetworkPy's methods in identifying physiologically meaningful metabolic changes, we present a series of case study analyses featuring data from the bacterial pathogen, *Mycobacterium tuberculosis* (Mtb). We used the Mtb GSMM iEK1011_v2 developed by [38] as the base model for these analyses. We report on two types of case studies, described below.

First, we compared genome-scale MetworkPy analysis outputs with genome-scale phenotype profiling data (See **S3.1**). The input phenotype profiling data we used were functional genomic screening data profiling gene essentiality (assessing whether disrupting a gene causes a growth or survival defect, measured by [39]; data retrieved from [40]) and gene vulnerability (assessing the extent of gene depletion that is required to elicit a growth or survival defect; vulnerable genes elicit defects with only partial depletion, measured by [40]). The goal of these comparisons was to show how network properties calculated by MetworkPy methods correlate with meaningful bacterial growth and survival outcomes.

Second, we assessed MetworkPy analysis outputs generated from data profiling the well-characterized Mtb transcription factor (TF), ArgR, which is known to regulate arginine biosynthesis [41, 42] (see **S3.1 – S3.4**). The input data for our analyses were published transcriptome profiles measured from a recombinant Mtb strain that induces ArgR overexpression [43]. We used these transcriptome profiles in two ways: 1) we defined genes that are significantly differentially expressed in these profiles as the set of regulatory target genes of ArgR, and we applied this target gene set to MetworkPy analyses; 2) we used the ArgR transcriptome profiles to generate iMAT-constrained condition-specific GSMMs that serve as inputs to MetworkPy analyses. The goal of these analyses was to show how MetworkPy methods can highlight metabolic changes that we expect to occur, since metabolic changes related to arginine metabolism should be highlighted with MetworkPy analyses of ArgR data. The results from our ArgR analyses are visualized on Escher maps of Mtb nitrogen and central carbon metabolism.

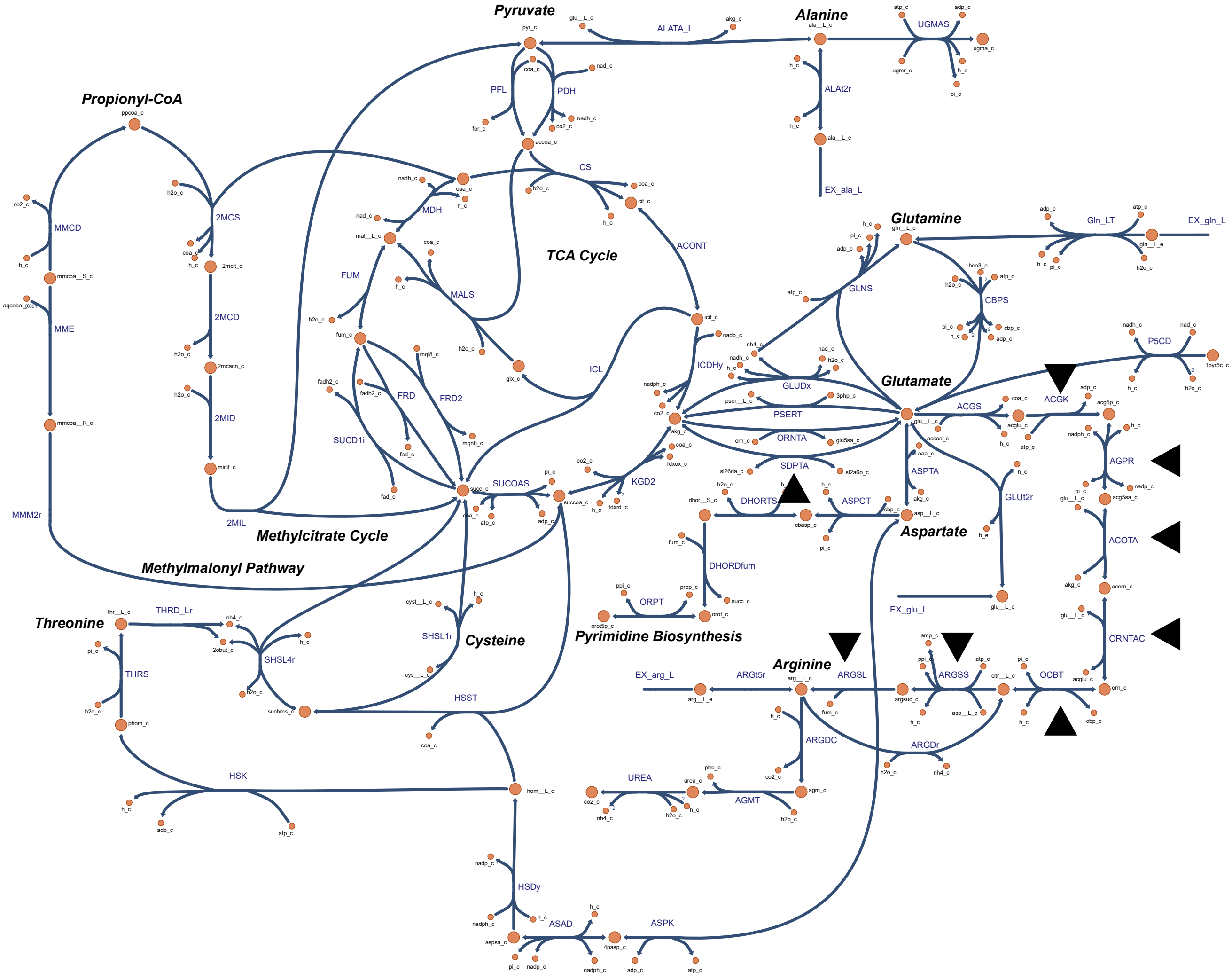


**Figure S11** Escher Map [17] of *Mycobacterium tuberculosis* nitrogen and central carbon metabolism. Both large and small circles indicate metabolites and the connecting curved or straight lines indicate reactions. Black triangles indicate the reactions associated with genes targeted by ArgR based on data from [43].

**S3.1 Target KO-Divergence**

To investigate the impact of all single-gene disruptions on Mtb metabolism, we performed KO-divergence analysis on the iEK1011_v2 model, *in silico*-disrupting each gene individually and evaluating the KL divergence between the flux distributions from resulting KO-model versus the flux distributions from the base iEK1011_v2 model. The results of this KO divergence analysis for the BIOMASS__2 reaction and the metabolite synthesis subnetworks are reported in the 'Gene KO Divergence' sheet of **Additional File 3**.

To examine the biological significance of the KO divergence values, we evaluated the association between the BIOMASS__2 reaction divergence in iEK1011_v2 and experimentally defined essential and vulnerable genes in Mtb [39, 40] Essential genes caused a significantly higher divergence than non-essential genes (Mann-Whitney U-test p = $4.5*10^{-66}$). Further, the BIOMASS__2 reaction divergence could predict gene essentiality with an AUC-ROC = 0.84. The biomass divergence is also significantly correlated with the vulnerability index from [40] (Pearson R = -0.61; p = $1.69*10^{-103}$).

As a point of comparison, we also performed single gene KO analysis using the traditional FBA approach with COBRApy. We found that the FBA-predicted biomass growth predictions correlated extremely closely with MetworkPy's biomass divergence (Pearson's R = -0.996). The AUC-ROC of predicting gene essentiality using the FBA KO method is identical to the AUC ROC of MetworkPy's divergence approach.

Beyond evaluating the impact of gene disruption on biomass production, MetworkPy's KO-divergence approach is also able to simultaneously evaluate the impact of single or multi-gene disruptions on the rest of metabolism. To explore the broader metabolic impacts of gene disruptions, we evaluated the divergence for metabolite synthesis networks following gene KO. Since many metabolite synthesis networks require similar sets of reactions, we grouped the metabolite synthesis networks to a representative set using correlation.

To group the metabolite synthesis networks, we calculated a pairwise correlation network, where the nodes represent metabolites and the edge weights represent the pairwise correlation between Boolean vectors representing whether a reaction is included in the synthesis subnetwork for those two metabolites. Edges were removed if they had a weight of less than 0.75. A connected component-based approach [44] identified groups of metabolites whose synthesis subnetworks were significantly correlated. From each group, we selected a single metabolite representative, based on which metabolite had the highest degree in the correlation network.

For each of these representative metabolites, we determined the KL divergence for their synthesis subnetwork between the single gene KO model and the iEK1011_v2 base model. We performed this calculation for each gene in the GSMM.

ArgR's regulated target genes had significantly higher KO-divergence for the representative metabolite associated with arginine, when compared against genes not targeted by ArgR (Benjamini-Hochberg adjusted Mann-Whitney U-test p = $5.22*10^{-4}$). ArgR regulatory target genes were defined based on data from [43], with targets being genes with a differential expression of at least a 2-fold magnitude upon TF overexpression relative to the median expression value, and an adjusted p-value of less than 0.05. The results of the analysis of the KO divergence for the gene regulatory targets of each of the Mtb TFs can be found in sheet 'TF Target KO Divergence' in **Additional File 3**.

**S3.2 ArgR Target Density and Enrichment**

To evaluate the metabolic enrichment of the regulatory target genes of ArgR, we generated a reaction stoichiometric connectivity graph from iEK1011_v2 and calculated the density and enrichment for reactions associated with the ArgR regulatory target genes. ArgR regulatory target genes were defined by the same criteria as in **S3.1** based on data from [43] (targets were genes with differential expression magnitude of at least 2-fold on TF overexpression relative to the median expression value, and an adjusted p-value of less than 0.05).

We evaluated the enrichment of these ArgR target genes in these reaction neighborhoods, as well as in the metabolite synthesis subnetworks. The results of this analysis can be found in the 'ArgR Rxn Neighbor Enrichment' sheet of **Additional File 3**. Two reactions showed significant enrichment of ArgR regulatory targets in their neighborhoods following false discovery (FDR) correction (Benjamini-Hochberg adjusted p-value<0.05): acetylglutamate kinase (ACGK, adjusted p = $9.1*10^{-4}$) and ornithine carbamyltransferase (OCBT, adjusted p = $1.8*10^{-2}$). Both reactions are in the "Arginine and Proline Metabolism" subsystem of iEK1011_v2. The significance of the reaction neighborhood enrichment in in Mtb's nitrogen metabolism is shown in **Figure S12B**. For the metabolite synthesis subnetworks, we found that the arginine biosynthesis network is significantly enriched for targets of ArgR (adjusted p = $1.86*10^{-11}$), as expected. The full metabolite network enrichment results of ArgR target in iEK1011_v2 can be found in the 'TF Metabolite Enrichment' sheet of **Additional File 3**.

We additionally evaluated the target density metric for ArgR's target genes mapped to Mtb*'s* nitrogen metabolism is shown in **Figure S12A**. The density metric shows that ArgR's target genes include several genes associated with reactions within the arginine biosynthetic pathway, as expected. Notably, the density metric's propagation through the network highlights how ArgR's target genes are also proximate to reactions outside of arginine biosynthesis. For example, 'ASPCT' (Aspartate carbamoyltransferase), a reaction in the "Purine and Pyrimidine Biosynthesis" subsystem of iEK1011_v2, has the 7$^{th}$ highest neighborhood density. Although ASPCT is not a direct member of "Arginine Biosynthesis" pathway, it is proximate to the reactions associated with arginine biosynthesis that are targeted by ArgR (**Figure S12A**). This network proximity to ArgR target genes, highlighted by the density metric, informs the hypothesis that ASPCT reaction activity (as well as its corresponding subsystem: Purine and Pyrimidine Biosynthesis) is likely to be impacted by ArgR overexpression. The density values for all the reactions in the iEK1011_v2 model can be found in the 'ArgR Rxn Neighborhood Density' sheet of **Additional File 3**.

As a comparison, we evaluated the hypergeometric enrichment of ArgR target genes in metabolic subsystems (defined in iEK1011_v2) and in KEGG pathways [45] (see 'TF Subsystem Enrichment' and 'TF KEGG Enrichment' sheets in **Additional File 3**). As expected, the "Arginine and Proline Metabolism" subsystem shows significant enrichment (adjusted p = $3.7*10^{-8}$). For the KEGG pathways, several pathways show significant enrichment: "Arginine biosynthesis" (adjusted p = $6.0*10^{-11}$), "Biosynthesis of amino acids" (adjusted p=$4.1*10^{-6}$), "2-Oxocarboxylic acid metabolism" (adjusted p=$1.0*10^{-3}$), and "Biosynthesis of secondary metabolites" (adjusted p=$1.2*10^{-2}$). Though this pathway enrichment approach is able to identify arginine biosynthesis (and pathways which subsume it), reactions and genes that fall outside of these pathway definitions but are nevertheless proximate are not easy to identify. For example, evaluating hypergeometric enrichment with "Purine and Pyrimidine Biosynthesis" subsystem discussed above showed no enrichment (p = 1.0) as there is no direct overlap between genes in that subsystem with ArgR targets. Similarly, evaluating the KEGG pathways "Purine metabolism" and "Pyrimidine metabolism" also yielded an enrichment p-value of 1.

We performed further analysis of the metabolite synthesis and consuming subnetworks using gene set variation analysis (GSVA) [29] (implemented in decoupleR [46]). To do this, we used the $\log_2$ fold-change expression data for the TF overexpression strains from [43], with MetworkPy-defined metabolite synthesis and consuming networks from iEK1011_v2 as the gene sets for GSVA. The metabolite networks with the highest GSVA statistic for ArgR were arginine and closely related metabolites, including Ornithine, N2-succinyl-L-gluatamate, N2-Succinyl-L-glutamate 5-semialdehyde, N2-Succinyl-L-ornithine, N2-Succinyl-L-arginine, N2-Succinyl-L-arginine, Agmatine, and L-Citrulline. All these metabolites are involved in, or proximate to, the biosynthesis of arginine. The results of GSVA can be found in the 'TF Metabolite GSVA' sheet of **Additional File 3**.

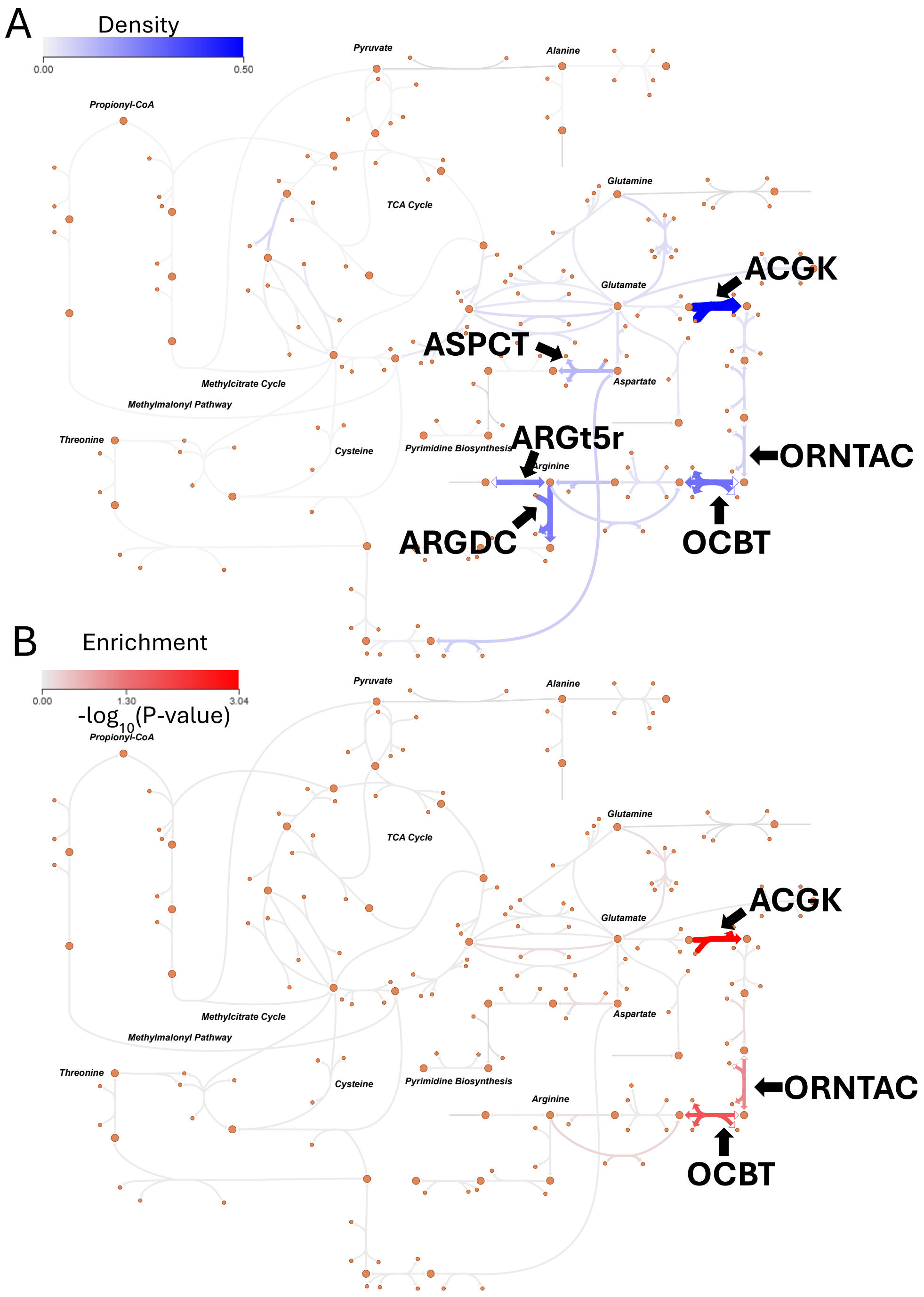


**Figure S12** Reaction neighborhood density and enrichment of ArgR regulatory targets visualized using the Escher web tool [17]. (A) Shows the reaction neighborhood density. The edges are labelled, colored, and sized according to the density of ArgR regulatory gene targets in a neighborhood, of radius 1, around the associated reaction. Higher density is shown in blue. (B) Shows the reaction neighborhood enrichment significance, equal to $-log_{10}(p - value)$. The edges are labelled colored and sized according to the enrichment of ArgR regulatory gene targets in a neighborhood, of radius 1, around the associated reaction. More significant enrichment (p-value closer to 0.0) is shown as red.

#### S3.3 Network Centrality

To investigate the parts of the metabolic network that ArgR regulates, we also applied the subset centrality metrics discussed in **S2.2.3** to the stoichiometric connectivity networks generated from iEK1011_v2.

We evaluated the centrality of nodes in the stoichiometric connectivity network with reference to the targets of ArgR. The full results of the subset centrality measures are included in sheet 'TF Target Subset Betweenness' (in **Additional File 3**).

The 5 reactions with the highest subset betweenness, after subtracting off the full network betweenness, are: ORNTAC (Ornithine transacetylase), AGPR (N-acetyl-g-glutamyl-phosphate reductase), ARGSS (Argininosuccinate synthase), ACOTA (Acetylornithine transaminase), and ASPTA (Aspartate transaminase), all of which are in the 'Arginine and Proline Metabolism' subsystem, or the closely related 'Alanine, Aspartate, and Glutamate Metabolism' (in **Additional File 3** see the 'TF Target Subset Betweenness' sheet for the full subset betweenness results, and see the 'Reaction Information' sheet for more details about these reactions, including their equations).

For the subset closeness centrality, the 5 reactions with the highest centrality are (after subtracting off the full network closeness): ACOTA (Acetylornithine transaminase), ORNTAC (Ornithine transacetylase), GSEALD (GSEALD), ARGSS (Argininosuccinate synthase), and OCBT (Ornithine carbamoyltransferase). ACOTA, ORNTAC, and OCBT are in the 'Arginine biosynthesis' subsystem, ARGSS is in the nearby 'Alanine, Aspartate, and Glutamate Metabolism', and GSEALD is in the 'Cofactor and Prosthetic Group Biosynthesis', which involved glutamate (in **Additional File 3** see the 'TF Target Subset Closeness' sheet for the full subset closeness results, and see the 'Reaction Information' sheet for more details about these reactions, including their equations).

The 5 metabolites with the highest subset betweenness centrality are: L-Glutamate, Ornithine, N-Acetyl-L-glutamate 5-semialdehyde, L-Aspartate, and N(omega)-(L-Arginino) succinate, all of which are associated with the 'Arginine biosynthesis' KEGG pathway. These highlighted metabolites overlap significantly with the 5 metabolites with the highest subset closeness centrality, which are: L-Glutamate, N2-Acetyl-L-ornithine, N-Acetyl-L-glutamate 5-semialdehyde, Ornithine, and L-Citrulline. All of which are again associated with the 'Arginine biosynthesis' KEGG pathway (see the 'Metabolite Information' sheet in **Additional File 3**).

#### S3.4 ArgR iMAT Flux Sampling Divergence Analysis (FSDA)

We evaluated the impact of ArgR on *Mtb* metabolism using transcriptome data profiling an ArgR overexpression strain [43]. We first used iMAT to generate expression-constrained GSMMs for each of the TF overexpression (TFOE) strains, with input transcriptomes from [43] and the iEK1011_v2 model from [38]. For each TFOE strain, we assigned genes with expression levels in the bottom 15% of that strain a weight of -1, and we assigned genes with expression levels in the top 15% (of that strain) a weight of 1 (all other genes received a weight of 0). These gene weights were then converted into reaction weights using the iEK1011_v2 model's GPR rules. We calculated the fluxes associated directly with the iMAT optimization problem and generated a model using the 'fva' iMAT approach described above. To generate a baseline model to compare against each of the TFOE iMAT models, we constructed an iMAT model using the median expression level of genes across the TFOE strains and assigned gene weights and reaction weights in the same way as for the TFOE strains.

We performed flux sampling for the TFOE and median models using the 'OptGP' sampler [47] implemented in COBRApy[3]. We then calculated the KL divergence of the flux distributions for reactions between each TFOE model and the median model, normalizing the divergence values across the TF models. **Figure S13** shows the normalized divergence values for the ArgR-overexpression condition-specific iMAT model on the nitrogen metabolism Escher map. The divergence highlights reactions throughout the arginine biosynthesis pathway (except for the OCBT reaction, which is caused by an iEK1011_v2 model artifact: iEK1011_v2 includes a loop that allows flux to bypass this reaction while still meeting iMAT restrictions). The genetic perturbation of ArgR overexpression also causes divergence in the biosynthetic pathways of other amino acids, including glutamate, glutamine, aspartate, and alanine. This prediction occurs due to the interconnected nature of the urea cycle and amino acid biosynthesis. The normalized divergence values for the reactions and metabolite synthesis networks for all the TFs can be found in the 'Normalized iMAT Reaction Div' and 'Normalized iMAT Metabolite Div' sheets of **Additional File 3** respectively. The reaction with the highest normalized divergence value for the ArgR model is ORNTA (Ornithine transaminase), in arginine and proline metabolism. Further, of the top 10 most divergent reactions, 6 are in the "Arginine and Proline Metabolism" subsystem, with the remainder in the closely related "Alanine, Aspartate, and Glutamate Metabolism". For the metabolite subnetwork divergence, the most divergent metabolite is ornithine (a precursor of arginine). Of the top 10 most divergent metabolites, 9 of them are associated with the "Arginine biosynthesis" or "Arginine and proline metabolism" KEGG [45] pathways (the other metabolite is isoleucine, which is connected to arginine metabolism through the ILETA reaction which converts between isoleucine and glutamate in the iEK1011_v2 model).

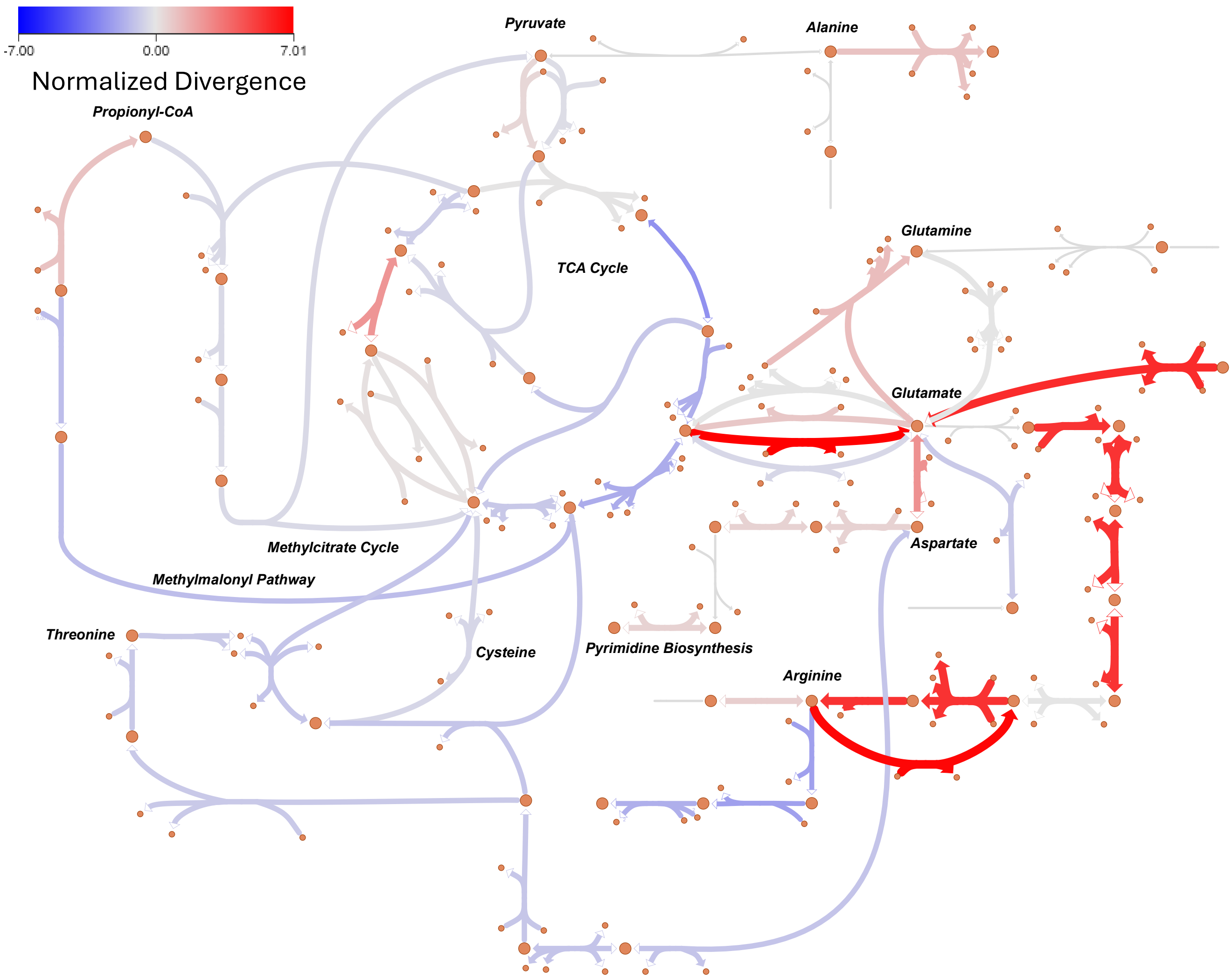


**Figure S13** Normalized flux divergence of ArgR iMAT model in nitrogen metabolism of Mtb visualized using the Escher web tool [17]. Flux divergence values were calculated for each TF between their iMAT constrained model and the median expression model. The divergence was then normalized across TFs to identify metabolic perturbations significantly associated with each specific TF. Edges are labelled, colored, and sized according to the divergence of the associated reaction, with more divergent reactions, compared to other TFs, shown in red and less divergent reactions, compared to other TFs, shown in blue.

To combine the information from FSDA and neighborhood density analyses, we used robust rank aggregation (RRA) [30]. Specifically, we used the non-normalized iMAT reaction divergence values to rank all the reactions in each TFOE condition-specific model and similarly ranked all the reactions based on their reaction neighborhood target density. These two rankings were then combined using the RRA algorithm to identify the top reactions associated with each TFOE strain (visualized in **Figure S14**, see the "RRA" sheet of **Additional File 3** for the full results).

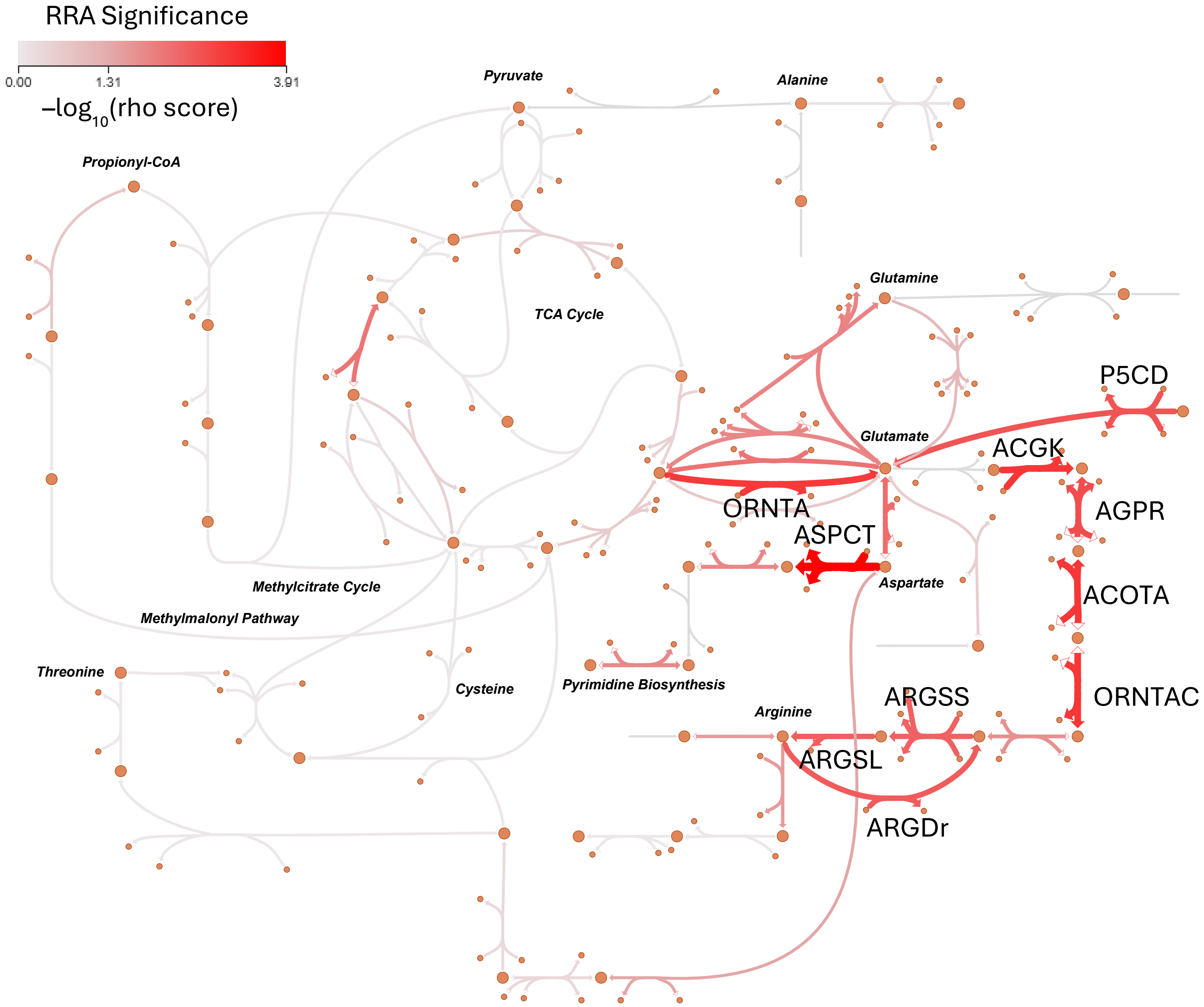


**Figure S14** Robust Rank Aggregation (RRA) combining target density and iMAT divergence results for ArgR in nitrogen metabolism of Mtb visualized using the Escher web tool [17]. Flux divergence was found for each TF between their iMAT constrained model and the median expression model. This was combined with the density of ArgR regulatory gene targets in a neighborhood, of radius 1, around the associated reaction using the Robust Rank Aggregation approach [30]. Reactions are colored and sized by the significance, equal to $-log_{10}(rho)$, where rho is the significance score from RRA [30]. More significant RRA (rho closer to 0.0) are shown in red. The top 10 reactions are labelled with their reaction ID from the iEK1011_v2 model.

For the ArgR overexpression strain, the top 10 reactions were (marked in **Figure S14**): Aspartate carbamoyltransferase (ASPCT), Acetylglutamate kinase (ACGK), Acetylornithine transaminase (ACOTA), Ornithine transacetylase (ORNTAC), Ornithine transaminase (ORNTA), N-acetyl-g-glutamyl-phosphate reductase (AGPR), 1-pyrroline-5-carboxylate dehydrogenase (P5CD), Arginine deiminase (ARGDr), Argininosuccinate lyase (ARGSL), Argininosuccinate synthase (ARGSS) (see sheet 'Reaction Information' of

**Additional File 3** for more information on these reactions). Of these reactions, 6 are in the ‘Arginine and Proline Metabolism’ subsystem, and another 3 are in the closely related ‘Alanine, Aspartate, and Glutamate Metabolism’ metabolism, with the remaining reaction being a consumer of aspartate in the ‘Purine and Pyrimidine Biosynthesis’ subsystem.

As a comparison analysis strategy, we found the parsimonious Flux Balance Analysis (pFBA) [13] solution using COBRApy for the iEK1011_v2 base model, the constrained ArgR iMAT model, and the iMAT model constructed using the median expression across the TFOE strains.

Examining the reactions which show the greatest difference between the unconstrained model pFBA and the ArgR FVA iMAT model, the top 10 reactions are in the “Redox Metabolism”, Oxidative Phosphorylation”, “Transport”, “Extracellular exchange” and “Citric Acid Cycle” subsystems. Similarly comparing the ArgR iMAT model to the median iMAT model, the reactions with the largest difference between the pFBA solutions are in the “Transport”, “Oxidative Phosphorylation”, “Redox Metabolism”, “Extracellular exchange”, “Purine and Pyrimidine Biosynthesis”, and “Nucleotide Salvage Pathway” subsystems. The full results of this pFBA-based iMAT analysis can be found in the “ArgR iMAT Fluxes” sheet of **Additional File 3**.

We also assessed various statistical methods of evaluating the differences in flux sample distributions between the ArgR iMAT model and the median iMAT model. We tested the Mann-Whitney U-test, Kolmogorov-Smirnov test, and the Kruskal-Wallis H-test (as implemented in SciPy [10]). The specific results of all these tests can be found in the “ArgR iMAT Fluxes” sheet of **Additional File 3**. In general, these methods identify most of the reactions in the model as being significantly different. Notably, it is difficult to differentiate the importance of many reactions as they have the same p-value (i.e. p = 0). For example, the t-test approach shows reactions in ‘Oxidative Phosphorylation,” “Transport,” and “Extracellular exchange” all having the same p-value (i.e. p = 0) as reactions in the “Arginine and Proline Metabolism” subsystem. The situation is the same for the Kolmogorov-Smirnov test, the Mann-Whitney U-test, and the Kruskall-Wallis H-test. Similarly, using these statistical tests to compare FVA iMAT versus unconstrained pFBA all do identify reactions in the “Arginine and Proline metabolism” subsystem as being significantly altered; importantly however, the tests cannot distinguish between arginine/proline subsystem reactions relative to other reactions across most of the rest of Mtb metabolism.
